# Acute Axon Damage and Demyelination are Mitigated by 4-Aminopyridine (4-AP) Therapy after Experimental Traumatic Brain Injury

**DOI:** 10.1101/2022.01.27.477989

**Authors:** Kryslaine L. Radomski, Xiaomei Zi, Fritz W. Lischka, Mark D. Noble, Zygmunt Galdzicki, Regina C. Armstrong

## Abstract

Damage to long axons in white matter tracts is a major pathology in closed head traumatic brain injury (TBI). Acute TBI treatments are needed that protect against axon damage and promote recovery of axon function to prevent long term symptoms and neurodegeneration. Our prior characterization of axon damage and demyelination after TBI led us to examine repurposing of 4-aminopyridine (4-AP), an FDA-approved inhibitor of voltage-gated potassium (Kv) channels. 4-AP is currently indicated to provide symptomatic relief for patients with chronic stage multiple sclerosis, which involves axon damage and demyelination. We tested clinically relevant dosage of 4-AP as an acute treatment for experimental TBI and found multiple benefits in corpus callosum axons. This randomized, controlled pre-clinical study focused on the first week after TBI, when axons are particularly vulnerable. 4-AP treatment initiated one day post-injury dramatically reduced axon damage detected by intra-axonal fluorescence accumulations in Thy1-YFP mice of both sexes. Detailed electron microscopy in C57BL/6 mice showed that 4-AP reduced pathological features of mitochondrial swelling, cytoskeletal disruption, and demyelination. Furthermore, 4-AP improved the molecular organization of axon nodal regions by restoring disrupted paranode domains and reducing Kv1.2 channel dispersion. 4-AP treatment did not resolve deficits in action potential conduction across the corpus callosum, based on *ex vivo* electrophysiological recordings at 7 days post-TBI. Thus, this first study of repurposing 4-AP as an acute treatment for TBI demonstrates pre-clinical efficacy in decreasing pathological hallmarks of axon damage. Studies beyond this acute phase are now warranted to assess functional utility and outcome trajectory.

**SIGNIFICANCE STATEMENT:** Traumatic brain injury (TBI) is an acute injury that, if unresolved, can progress to cause persistent, debilitating symptoms. Currently, no treatments effectively prevent damage to long myelinated axons in white matter tracts, which is a hallmark pathology of TBI. 4-aminopyridine (4-AP) is FDA-approved to treat chronic symptoms in patients with multiple sclerosis, which involves autoimmune damage to myelinated axons. As the first assessment of repurposing 4-AP as an acute treatment for TBI, our randomized, controlled studies tested the hypothesis that low-dose 4-AP initiated one day after experimental TBI will reduce acute axon damage and demyelination. We found that 4-AP treatment significantly reduced the progression of axon pathology and demyelination during the first week after TBI using clinically relevant experimental conditions.

## INTRODUCTION

Traumatic brain injury (TBI) results in long term disability in moderate to severe cases and can cause persistent debilitating symptoms even in patients with a “mild” diagnosis (Juengst et al., 2020; Levin et al., 2021; McCrea et al., 2021; Yue et al., 2021). Traumatic injury to long axons within white matter tracts is a key pathology in all forms of closed head TBI. Importantly, white matter injury represents a treatable target for TBI and neurodegenerative diseases, including multiple sclerosis (MS) and Alzheimer’s disease (Filley and Kelly, 2018; Simon and Watkins, 2018; Andravizou et al., 2019; Roseborough et al., 2020).

The lack of treatments for TBI, particularly treatments to protect axons that do not effectively regenerate, is a critical gap in research and clinical care. At an early stage of damage, axons have the potential to recover but once mechanical or molecular processes fragment the axon, then irreversible axon degeneration proceeds (Marmarou et al., 2005; Nikic et al., 2011; Maxwell et al., 2014; Williams et al., 2014; Gu et al., 2017; Loring and Thompson, 2020). Additionally, axons that remain viable can undergo demyelination, i.e. loss of their myelin sheaths, which impairs rapid signal conduction and desynchronizes neural circuits. Axon damage and demyelination contribute to slow information processing and neural circuit deficits that often underlie symptoms of mild-moderate TBI (Dams-O’Connor et al., 2013; Donders and Strong, 2014; Dymowski et al., 2016). Maintaining axon and myelin function can alleviate symptoms and prevent irreversible white matter degeneration after TBI. No available treatments effectively protect against axon damage or promote remyelination, which protects axons and enhances recovery (Pan and Chan, 2017; Koliatsos and Alexandris, 2019).

Treatment strategies for TBI, which have failed to date, have not focused on axon or myelin damage in the context of white matter pathology. Our studies of white matter injury revealed axon and myelin damage that included disruption of nodal regions and slowed conduction velocity early after TBI (Mierzwa et al., 2015; Marion et al., 2018). These findings resemble aspects for which 4-aminopyridine (4-AP) was developed to treat patients with MS. 4-AP is prescribed for patients in the chronic stage of MS and responders have positive effects on walking ability, finger dexterity, and cognitive function (Zhang et al., 2021). Recent pre-clinical testing on acute peripheral nerve injuries, which also involve axon and myelin damage, revealed the surprising result that 4-AP treatment enhances structural recovery of axons and myelin (Tseng et al., 2016). Thus, 4-AP may be beneficial for acute traumatic injury and warrants evaluation of repurposing for TBI.

4-AP is a small molecule inhibitor of voltage-gated potassium (Kv) channels that readily crosses the blood-brain barrier yet has an uncertain mode of action in patients (Goodman and Stone, 2013; Dietrich et al., 2021). In MS and other chronic neurological diseases, 4-AP is thought to enhance action potential conduction and modulate overall neural circuit activity through inhibiting aberrant Kv currents of axons and/or synapses (Smith et al., 2000; De Giglio et al., 2020). Therapeutically, low dose (10 mg, twice daily) extended release 4-AP (dalfampridine) is effective at blood levels below 100 ng/ml (∼ 1 μM) without unacceptable adverse effects, particularly seizure induction (Goodman and Stone, 2013).

We conducted randomized, controlled pre-clinical studies to evaluate repurposing of 4-AP as an acute treatment for TBI. Low therapeutically relevant 4-AP dosing was initiated one day after TBI to evaluate axon integrity and electrophysiological function during the first week, when axons are particularly vulnerable. A single impact closed head TBI in adult mice produced traumatic axonal injury in the corpus callosum (CC). This injury models human TBI pathology and neuroimaging features from acute injury through chronic neurodegeneration (Sullivan et al., 2013; Mierzwa et al., 2015; Marion et al., 2018; Bradshaw et al., 2021b). The current study design demonstrates reproducible CC pathology and functional deficits to rigorously evaluate acute effects of 4-AP treatment. We identify multiple positive effects of 4-AP treatment on mitigating axon pathology after TBI.

## MATERIALS AND METHODS

### Study Design

Experiments in this study (Figure 1) were performed using a blinded, randomized, controlled trial design (Henderson et al., 2013). Animal procedures were designed and implemented according to the ARRIVE (Animal Research: Reporting In Vivo Experiments) guidelines (Kilkenny et al., 2010). Pre-determined study designs with details for data collection, standardized method protocols, outcome measures, sample size estimation (> 80% power), statistical analyses, and rules for data exclusion were pre-registered on Open Science Framework (https://osf.io/registries) prior to executing experiments. Each investigation followed a pre-determined study design to evaluate 4-AP treatment using three complementary approaches: 1) screen for axon damage and analyze the molecular organization at nodes of Ranvier by confocal microscopy using Thy1-YFP-16 reporter mice; 2) conduct in-depth axon-myelin pathology by electron microscopy (EM) using C57BL/6J mice; and 3) directly examine axon function by *ex vivo* electrophysiological recordings using coronal brain slices from C57BL/6J mice.

**Figure 1.**
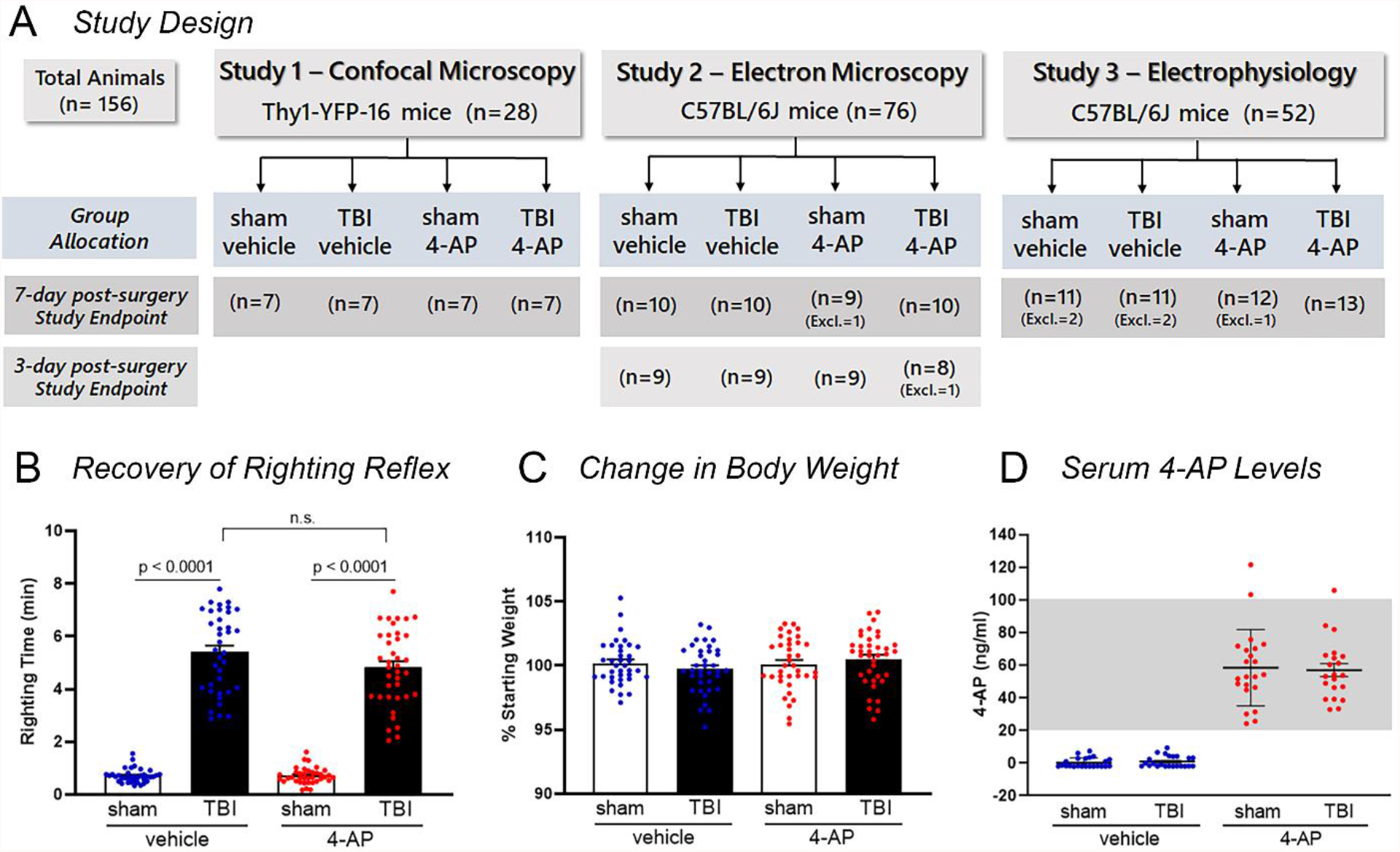
Experimental study design and key assessments of group allocation. (A) Graphical representation of experimental timeline and treatment groups. Three sets of experiments were conducted according to pre-determined study designs. For each study, the strain and number of mice used is shown along with the study endpoint and method of analysis of corpus callosum axons. Excl. = number of mice excluded based on pre-determined criteria (see Materials and Methods for definition). (B) TBI increased the time to right from supine to prone position after surgery (righting reflex) as compared with respective sham groups. Mice randomized to the TBI vehicle and TBI 4-AP groups (before drug administration) exhibited similar righting times. (C) 4-AP treatment did not cause a significant change in body weight over the period of the experiment. (D) 4-AP serum drug level concentrations are within the clinical therapeutic target range (20-100 ng/ml) in cardiac blood collected approximately 30-60 min after last 4-AP injection. Graphs show the mean ± SEM with each mouse as an individual data point. One-way ANOVA with Tukey’s multiple comparisons test: (B) F_3,145_ = 211.3, p < 0.0001. Two-way ANOVA with Holm-Sidak *post hoc* test: (C) Interaction: F_1,145_ = 1.936, p = 0.1663, Injury: F_1,145_ = 0.009971, p = 0.9206, Drug: F_1,145_ = 1.209, p = 0.2733. Cohen’s *d* effect size: (B) sham veh vs TBI veh: *d* = 4.322, sham 4-AP vs TBI 4-AP: *d* = 3.919.

### Randomization and Blinding Procedures

All animals were subcutaneously implanted with a micro-transponder (IMI-500, Bio Medic Data Systems, Seaford, DE) two days before surgery and the designated alpha-numeric codes were used for randomization and animal group allocation prior to surgical procedures. Computer-generated randomization was achieved by first assigning mice to TBI or sham surgical groups, then to vehicle or 4-AP treatment groups using the RANDBETWEEN function in Microsoft Excel (RRID:SCR_016137). Both the investigators administering the drug/vehicle and those performing data analyses were blinded as to injury and treatment conditions. Unblinding only occurred after all data were collected and statistical analyses were completed. The study designs, including the number of animals randomized to sham-vehicle, TBI-vehicle, sham-4-AP, or TBI-4-AP groups with endpoints at either 3- or 7-days post-injury are shown in Figure 1A.

### Mice

All animal experiments were carried out in accordance with the National Institutes of Health Guide for the Care and Use of Laboratory Animals and in compliance with protocols approved by the Institutional Animal Care and Use Committee at the Uniformed Services University of the Health Sciences. At the start of experiments, mice were 8-10 weeks old, which approximately corresponds to young adulthood in humans (Dutta and Sengupta, 2016), a patient population at high risk of TBI (Taylor et al., 2017). Transgenic mice expressing YFP under the control of the Thy-1 promoter (Thy1-YFP-16) on a C57BL/6 background were acquired from Jackson Laboratories (Bar Harbor, ME). The original Thy1-YFP-16 mice (IMSR Cat# JAX:003709, RRID:IMSR_JAX:003709; B6.Cg-Tg(Thy1-YFP)16Jrs/J) were bred in-house to generate the experimental male (n=3/group) and female (n=4/group) mice for Study 1 – Confocal Analysis. As no sex-based differences were detected in this initial study, the experiments of Studies 2 and 3 used only male mice. C57BL/6J (IMSR Cat# JAX:000664, RRID:IMSR_JAX:000664) mice were purchased from Jackson Laboratories. Mice were socially housed in 35 cm x 16.5 cm x 18 cm enrichment cages (2-5 mice per cage), exposed to 12:12 h light-dark cycle, at controlled room temperature, and with free access to food and water. All experimental procedures were conducted during the daytime light cycle (0600-1800) after the mice had acclimated for at least 3 days in their home cage.

### TBI Model

A single impact closed head TBI model was used to produce traumatic axonal injury in the CC that models traumatic axonal injury in human TBI pathophysiology (Sullivan et al., 2013; Mierzwa et al., 2014; Mierzwa et al., 2015; Yu et al., 2017; Marion et al., 2018; Bradshaw et al., 2021b). An Impact One Stereotaxic Impactor (Leica Biosystems, Buffalo Grove, IL) attached to a steel impactor with a 3-mm flat tip with rounded edges (Cat# 2520-3S, Neuroscience Tools, Aurora, IL) was zeroed on the surface of the skull centered at bregma (0 ML, 0 AP, 0 DV). In the coronal plane, bregma is aligned with the mid-line crossing of the anterior commissure, which served to localize the injury site in coronal brain sections. For mice in the TBI groups, the impactor was set to 4 m/s with a dwell time of 0.1 s and a depth of 1.5 mm from the surface of the skull (0 ML, 0 AP, -1.5 DV). Isoflurane administration was terminated immediately prior to impact. Our prior studies have shown that these parameters induce mild-moderate closed skull TBI with CC pathology that progresses to late stage CC atrophy without gross tissue damage or microhemorrhages, detected by neuropathology or magnetic resonance imaging (Mierzwa et al., 2015; Yu et al., 2017; Marion et al., 2018; Bradshaw et al., 2021b). Sham mice followed identical procedures but did not receive the impact. After scalp closure, mice were placed in a warm cage in the supine position to record the righting reflex as a measure of the time it takes each mouse to regain consciousness. A delay in the motor ability for a mouse to right itself from a supine position (righting reflex) immediately after the surgical procedure indicates a transient alteration of consciousness that is increased after TBI (Morehead et al., 1994; Dewitt et al., 2013).

### 4-Aminopyridine Preparation and Dosing

4-aminopyridine (4-AP, fampridine; Cat# 275775; ≥ 99% purity; Sigma-Aldrich, St. Louis, MO) was dissolved in sterile saline (0.9% sodium chloride; Hanna Pharmaceutical Supply Co., Inc.; Cat# 0409488810) to a final concentration of 0.1 µg/µl, and intraperitoneally (i.p.) injected at a dose of 0.5 mg/kg body weight. 4-AP dosing was calculated to approximate a relevant human clinical dose equivalent (clinical serum dose range of 20-100 ng/ml) as reported in prior rodent studies that have demonstrated 4-AP positive effects after peripheral nerve and pyramidal tract injuries (Tseng et al., 2016; Sindhurakar et al., 2017). Mice were injected i.p. twice a day (11-13 hrs apart) with either 4-AP solution or equivalent volumes of saline. Mice were weighed at baseline (surgery day) and on each day to determine the appropriate dose. Injections were initiated at 24 hours after sham or TBI procedures to align with a clinically reasonable interval for the time to first dose. The study endpoint was either 3- or 7-days post-sham or TBI procedures. For *ex vivo* electrophysiological studies, mice did not receive a final dose on day 7 post-injury in order to examine the endogenous electrophysiology without potential variation due to the *in vivo* 4-AP levels at the time of brain collection. Mice did not exhibit overt changes in coat, posture, or activity with 4-AP treatment. The percentage change in body weight at the study endpoint was calculated relative to each animal’s baseline as a measure of overall health relative to 4-AP treatment or TBI.

### 4-AP Quantification in Serum

For analysis of 4-AP serum levels, mice received the last 4-AP or vehicle injection approximately 30-60 min before cardiac blood was collected under deep anesthesia just prior to transcardial perfusion. Blood was undisturbed at room temperature for a 1 hr clotting time, and then serum was separated by centrifugation at 3,000 x g for 10 min and stored at -80°C. 4-AP serum concentration was analyzed by liquid chromatography coupled with tandem mass spectrometry (LC-MS/MS) as previously detailed (Hsu et al., 2020). All data points are included in the analyses except for: 1) one cohort (n = 3/group) from the 3-day C57BL/6J mice (Study 2), which had blood collected at longer times after injection for analysis of drug washout; 2) one mouse from the 7-day C57BL/6J cohort (Study 2) for which we were unable to collect enough blood; and 3) all mice used for electrophysiological recordings since the experiment was designed with a one day washout period in order to record the endogenous electrophysiological function.

### Confocal Microscopy

Thy1-YFP-16 reporter male and female mice were perfused with 4% paraformaldehyde (PFA). Collected brains were post-fixed in the same fixative overnight, cryoprotected and then cut as 14 µm coronal cryostat sections. The region-of-interest (ROI) was located in equivalent areas of the medial CC situated between the cingulum and the longitudinal fissure in coronal brain slices under the impact site. High resolution images were acquired on a Zeiss LSM 700 confocal microscope (RRID:SCR_017377) using either a 40x/1.4 or a 63x/1.4 oil Plan-Apochromat objective, for analyses of YFP+ accumulations and nodal domains respectively.

### Confocal Analysis of Axonal Swellings in Thy1-YFP-16 Mice

For each mouse, three separate confocal image stacks were analyzed and averaged to obtain a single value per mouse. The confocal image stacks were acquired with a volume of (mean ± SD) 219.8 ± 5.6 µm × 98.7 ± 13.0 µm × 4.90 µm (14 optical slices per stack) centered over the medial CC in either cerebral hemisphere. After conversion of image z-stacks to maximum intensity projections, images were processed using ImageJ software (NIH, RRID:SCR_003070) for conversion to grayscale and spatial calibration. In order to generate a binary mask for automated particle analysis, an adequate threshold was selected for each image based on an optimal separation of gray levels in swollen axons exhibiting strong YFP+ accumulations against background that included intact YFP-expressing axons. The ImageJ particle analysis plug-in was then used for quantification over the entire image (excluding edges). This process enabled automated quantification of the percent area containing YFP accumulations within the defined imaged volume. Results were analyzed for sex as a biological variable and as no significant differences were found, the data from males and females were combined.

### Confocal Analysis of Nodal and Paranodal Domains

Immunohistochemistry, confocal imaging, and quantification of paranodal complexes were described in detail previously (Marion et al., 2018). Briefly, detection of Nav1.6 (Alomone Labs, Jerusalem, Israel; Cat# ASC-009, RRID:AB_2040202) at the nodes of Ranvier and Caspr (UC Davis/NIH NeuroMab Facility Cat# K65/35, RRID:AB_2877274) at adjacent paranodes. Nav1.6 and Caspr localization was detected on confocal images acquired with a volume of 67.74 µm × 67.74 µm × 4.98 µm using a 0.45 µm z-interval separation between image planes. For each mouse, one image stack per hemisphere was collected and analyzed. Paranode domain organization was assessed by manual length measurements of 100 randomly selected pairs of Caspr domains for each cerebral hemisphere (total of 200 Caspr domains per mouse) in the 2D plane using maximum intensity z-projections (MIP). The average length difference within paranode pairs (paranodal asymmetry), the nodal gap length (distance between paired Caspr immunolabeled domains), and the total number of heminodes (unpaired Caspr domain adjacent to Nav1.6 immunolabeled node region) were tabulated. Because YFP fluorescence in TBI-induced axonal swellings can readily differentiate injured from non-injured tissue sections from Thy1-YFP-16 mice, the initial analysis did not include the YFP channel in order to avoid potential bias. After quantification was completed, confocal z-stacks were re-assessed using the 3-dimensional viewing software arivis Vision 4D (RRID:SCR_018000) to confirm that the identified unpaired Caspr domains (heminodes) met the pre-determined criteria of expression within YFP-expressing axons as detailed in Marion et al., 2018. Only Caspr domains localized along YFP axons were included in the analysis.

### Confocal Localization of Juxtaparanodal Kv1.2 Channel Domain

Cryostat sections from Thy1-YFP-16 mice were subjected to antigen retrieval with Tris-EDTA buffer (pH 9.0) in a 90°C water bath for 15 min. Co-labeling with primary antibodies for the juxtaparanode marker Kv1.2 (UC Davis/NIH NeuroMab Facility Cat# K14/16, RRID:AB_2877295) and the paranode marker Caspr (Abcam, Cambridge, UK; Cat# ab34151, RRID:AB_869934) was conducted overnight at 4°C. Sections were then exposed overnight to secondary antibodies donkey anti-mouse Alexa Fluor 594 (Cat# 715-586-151, RRID:AB_2340858) and donkey anti-rabbit Alexa Fluor 647 (Cat# 711-606-152, RRID:AB_2340625) both from Jackson ImmunoResearch (West Grove, PA). Kv1.2 and Caspr localization was analyzed on confocal image stacks of 11 µm in depth and measuring (mean ± SD) 131.0 ± 66.2 µm × 13.6 ± 15.9 µm in an area over the medial CC. Given the sensitivity of YFP expression to axon damage, Kv1.2 and Caspr domains were first analyzed without the YFP channel visible in order to maintain the investigator blinded to surgical group allocation. ImageJ Cell Counter plugin was used to classify all identifiable Kv1.2 immunolabeled juxtaparanodal domains within the MIP image as either “typical” or “atypical”. Typical Kv1.2 expression did not overlap with Caspr immunolabeled paranodal domains and appeared symmetrical in length. Atypical Kv1.2 expression exhibited > 30% overlap with the Caspr domain and/or appeared asymmetrical, often with Kv1.2 dispersion away from the nodal region into the internode. After completion, each classified Kv1.2 nodal complex was reanalyzed by sequentially examining individual optical planes with the YFP signal on using the Zeiss Zen Black software (ZEISS, RRID:SCR_018163). Only Caspr/Kv1.2 domains contained within YFP+ axons were included in the analysis. The percentage of typical and atypical Kv1.2 immunoreactive domains for each mouse was calculated based on the corresponding image area analyzed.

### Electron Microscopy

Transmission EM was performed to assay the efficacy of 4-AP treatment by detecting ultrastructural pathology, including myelin integrity, of individual axons as in prior studies (Mierzwa et al., 2015; Marion et al., 2019; Bradshaw et al., 2021b). C57BL/6J male mice were perfused with fixative containing 2.5% glutaraldehyde and 4% PFA in 0.1M phosphate buffer pH 7.4. The brains were dissected and post-fixed in the same fixative overnight. Sagittal 40 µm vibratome slices cut and osmicated to identify the CC region-of-interest (ROI), located above the lateral ventricles at the level of the crossing of the anterior commissure, for thin sectioning (∼70 nm) followed by processing for TEM.

### Quantification of Axon and Myelin Damage

To assess the extent of ultrastructural pathology, axons were classified by an investigator blinded to surgical and treatment allocation. Twenty-to-fifteen representative images from each mouse were collected at 5,000x magnification using a JEOL JEM-1011 TEM (JEOL USA Inc., Peabody, MA) and an AMT XR50S-A digital camera (Advanced Microscopy Techniques, Woburn, MA) from which 10 images were randomly selected for analysis using the Excel RANDBETWEEN function in order to minimize investigator bias. A 17.8 µm x 13.4 µm counting frame was drawn over each image using Adobe Photoshop (RRID:SCR_014199) and axons touching the left and bottom margins of the counting frame were excluded from quantification. Damaged axons exhibiting abnormal mitochondrial swelling (> 50% of transverse axon area) were classified separately from axons exhibiting damage based on neurofilament compaction or vesicle/organelle accumulation. Axons large enough to typically be myelinated (diameter > 0.3 µm for adult mouse CC; (Sturrock, 1980) but lacking myelin were classified as de/unmyelinated. Experiments were conducted as cohorts of 2-3 mice for each of the four treatment groups, with 3 cohorts combined for the 3-day endpoint and 4 cohorts for the 7-day endpoint to generate the mouse numbers shown in Figure 1A.

### Quantification of Axon Diameter, Myelin Thickness, and g-Ratio

For each mouse, a grid measuring 3 µm x 3 µm was laid over one of the randomly selected EM images described above. All myelinated axons (range = 61 to 120 axons) contained within or touching the perimeter of every 3^rd^ grid cell within the counting frame were measured, and then the first 50 axons meeting the circularity criteria were included in the statistical analysis. The freehand selection tool in ImageJ was used to trace the axonal circumference generating automatic calculations of axon area and circularity. Axon diameter was calculated from axon area values using the formula 2 x v(area/π). Myelin thickness was determined based on the average of radial line measures across the myelin sheath taken at four equidistant points around the axon (Zhou et al., 2012; Marion et al., 2019). Fiber diameter is 2 x myelin thickness + axon diameter. The g-ratio is the ratio of the axonal diameter to the fiber diameter. The fifty axons per mouse used in final calculations were approximately circular (having circularity > 0.6, where 1 = perfect circle) in order to avoid skewing of the data due to irregularly shaped axons or axons cut at a slant. In total, 350 axons were analyzed per sham animal group and 450 axons per TBI vehicle and TBI 4-AP groups.

### *Ex Vivo* Electrophysiology of CC Axons

Brain slices were used for *ex vivo* recordings to directly examine the electrophysiological properties of axons across the CC midline (Marion et al., 2018). Following isoflurane anesthesia, brains were collected and submerged in ice-cold artificial cerebrospinal fluid (ACSF) with sucrose (in mM: 2 KCl, 1 CaCl_2_, 1.25 NaH_2_PO_4_, 2 MgSO_4_, 2 MgCl-6H_2_O, 26 NaHCO_3_, 10 D-glucose, 206 Sucrose), bubbled with a gas mixture of 95% O_2_/5% CO_2_. Coronal sections of 400 µm thickness were sliced using a Leica VT1200 vibratome (Leica Biosystems Inc., Buffalo Grove, IL) containing sucrose ACSF. Sections were transferred to a holding chamber filled with normal ACSF (in mM: 126 NaCl, 3 KCl, 2 CaCl_2_, 1.25 NaHPO_4_, 2 MgSO_4_, 2 MgCl-6H_2_O, 26 NaHCO_3_, 10 D-glucose; bubbled with 95% O_2_/5% CO_2_ mixture) at 36°C for 30 min and then to room temperature for at least 1 hr prior to recording. Individual slices were then transferred to a perfusion and recording chamber (Warner Instruments, Hamden, CT) on an upright Zeiss Examiner Z1and perfused with normal ACSF continuously bubbled with 95% O_2_/5% CO_2_ at a rate of 1-2 ml/min at room temperature. A bipolar tungsten electrode (10k Ohm impedance) connected to a stimulus generator (S88K, Grass technologies, West Warwick, RI) was used for stimulation, while a borosilicate glass capillary (1M Ohm resistance in the bath) was used as the recording electrode. Evoked field potentials in the CC were recorded from axons near the recording electrode placed approximately 1.5 ± 0.18 mm from the stimulating electrode.

### Compound Action Potential (CAP) and Velocity Measurements

The evoked CAP was captured using a MultiClamp 700B amplifier (Molecular Devices). Data was acquired with pClamp10 (pClamp; Molecular Devices, RRID:SCR_011323) and analyzed with Clampfit 10.4 (Molecular Devices) and OriginPro 2020b (OriginLab, Northampton, MA) software packages. Parameters of stimulation protocols were as follows: (a) Input-output recording: 50 µA to 700 µA in 50 µA steps, 0.2 ms duration; (b) Velocity measurement: three additional positions for the recording pipette were placed successively closer to the stimulus electrode (stimulus strength adjusted accordingly). OriginLab Pro software (RRID:SCR_014212) was used to determine: (a) CAP amplitude, measured from the peak of the CAP to a tangent line drawn over the projected CAP bases; (b) CAP width, computed at the CAP half-height; and (c) CAP peak to recovery, the latency from the peak to the return of the positive phase of the CAP. Conduction velocity was calculated based on the best-fit linear regression analysis of distance between electrodes (ranging from 0.4 to 1.8 mm) versus recording of the time interval between CAP waveform peaks. Conduction velocity and CAP measures were acquired for all mice included in the analysis. After the pre-determined interim data analysis at n = 6 mice per condition, 7 more mice were added per condition (see Exclusion section below). After completing recordings for CAP and conduction velocity, the brain slices of the added mice were also tested for more specific conduction parameters of refractoriness and strength-duration testing, which were not included in the pre-determined study design.

### Refractoriness and Strength-Duration Analyses

For refractoriness testing, after a control single pulse was generated with 500 µA for 0.2 ms as stimulus, a paired pulse protocol (Reeves et al., 2005; Crawford et al., 2009) was implemented in which pulses of the same strength and duration were separated by an inter-pulse interval (IPI) ranging from 1.5 ms to 8 ms in 0.5 ms increments. The paired pulse protocol was then followed by another control single pulse run. The average of the two control run amplitudes was used as 100% single pulse value. The ratio of the amplitude of each CAP component after the second paired pulse stimulation to the amplitude of the corresponding CAP amplitudes from the single pulse stimulation were plotted as a function of the interpulse interval. The 50% ratio of CAP_2_/CAP_1_ was used to compare refractoriness across groups. For the Strength-duration measurement, after a control single pulse run (500 µA strength for 0.2 ms; considered 100% amplitude), recordings with increasing duration and adjusted stimulus strength were performed to elicit approximately 30% of the N1 and N2 amplitude of the control run. Duration steps were 50, 60, 80, 100, 120, 140, 160, 200, 300, 400, 500, and 600 ms. Finally, a final control run was recorded with 0.2 ms duration and 500 µA strength to account for changes of recording conditions over time.

### Statistical Analyses

Experimental data were analyzed and graphically represented using GraphPad Prism version 9.0.2 software (RRID:SCR_002798). Results are presented as mean ± standard error of the mean (SEM) unless otherwise noted. Two-group comparisons were made using unpaired Student’s *t*-test. One-way ANOVA tests were conducted when a single variable factor (injury) was tested in the analysis for the 4 animal groups (i.e., before 4-AP administration). Two-way ANOVA was used when interactions of both variable factors (injury and drug) were analyzed (see Tables 1-4 for results). When the ANOVA test was found to be statistically significant, *post hoc* Tukey or Holm-Sidak tests were used to make pairwise comparisons between group means. For evaluating compound action potential and strength-duration data, a linear mixed effects model was used with injury and drug as random effects and current intensity as fixed effect, followed by post hoc Holm-Sidak multiple comparisons test. Multiple counts derived from sampling in a given mouse were averaged and treated as one data point in each quantitative figure, except for the g-ratio x axon diameter plot which depicts all axons measured in the analysis, and the CAP, strength-duration and refractoriness data whose data points depict group means. Effect size was estimated using Cohen’s *d* formula as the mean difference between two groups divided by the pooled standard deviation. Cohen defined *d* = 0.2, *d* = 0.5 and *d* ≥ 0.8 as small, medium and large effects, respectively (Cohen, 1988). All statistical tests were two-tailed and significance was set as p < 0.05.

**Table 1.** Study 1 — Immunohistochemistry of axons in Thy1-YFP-16 mice (Figures 2 and 3). Student’s t test or Two-way ANOVA results with Holm-Sidak *post hoc* test and Cohen’s *d* effect size.

| Fig 2- Outcome Measures | t, df or F (DFn, DFd); p-value | Adjusted p-value | Cohen's *d* effect size |
| --- | --- | --- | --- |
| YFP Particles | $t_{12} = 9.785$; $p < 0.0001$<br><u>Interaction:</u><br>$F_{1,10} = 0.098$, $p = 0.7607$<br><u>Sex:</u><br>$F_{1,10} = 0.022$, $p = 0.8852$<br><u>Drug:</u><br>$F_{1,10} = 78.30$, $p < 0.0001$ | <u>Males:</u><br>TBI veh vs TBI 4-AP:<br>$p = 0.0006$<br><u>Females:</u><br>TBI veh vs TBI 4-AP:<br>$p = 0.0002$<br>4-AP: TBI males vs TBI females:<br>$p = 0.9381$ | TBI veh vs TBI 4-AP:<br>$d = 5.231$ |
| % Area of YFP Particles | $t_{12} = 10.05$; $p < 0.0001$<br><u>Interaction:</u><br>$F_{1,10} = 1.205$, $p = 0.2980$<br><u>Sex:</u><br>$F_{1,10} = 0.00000275$, $p = 0.9987$<br><u>Drug:</u><br>$F_{1,10} = 98.41$, $p < 0.0001$ | <u>Males:</u><br>TBI veh vs TBI 4-AP:<br>$p = 0.0005$<br><u>Females:</u><br>TBI veh vs TBI 4-AP:<br>$p < 0.0001$<br>4-AP: TBI males vs TBI females:<br>$p = 0.7028$ | TBI veh vs TBI 4-AP:<br>$d = 5.370$ |
| Fig 3- Outcome Measures | F (DFn, DFd); p-value | Adjusted p-value | Cohen's *d* effect size |
| Caspr+ Heminodes | <u>Interaction:</u><br>$F_{1,12} = 17.21$, $p = 0.0013$<br><u>Injury:</u><br>$F_{1,12} = 61.00$, $p < 0.0001$<br><u>Drug:</u><br>$F_{1,12} = 16.80$, $p = 0.0015$ | <u>sham veh vs TBI veh:</u><br>$p < 0.0001$<br><u>sham 4-AP vs TBI 4-AP:</u><br>ns, $p = 0.0651$<br><u>TBI veh vs TBI 4-AP:</u><br>$p = 0.0003$ | <u>sham veh vs TBI veh:</u><br>$d = 5.591$<br><u>sham 4-AP vs TBI 4-AP:</u><br>$d = 1.917$<br><u>TBI veh vs TBI 4-AP:</u><br>$d = 3.713$ |
| Caspr Length Difference | <u>Interaction:</u><br>$F_{1,12} = 15.66$, $p = 0.0019$<br><u>Injury:</u><br>$F_{1,12} = 31.46$, $p = 0.0001$<br><u>Drug:</u><br>$F_{1,12} = 15.66$, $p = 0.0019$ | <u>sham veh vs TBI veh:</u><br>$p = 0.0001$<br><u>sham 4-AP vs TBI 4-AP:</u><br>ns, $p = 0.6040$<br><u>TBI veh vs TBI 4-AP:</u><br>$p = 0.0005$ | <u>sham veh vs TBI veh:</u><br>$d = 5.340$<br><u>sham 4-AP vs TBI 4-AP:</u><br>$d = 0.754$<br><u>TBI veh vs TBI 4-AP:</u><br>$d = 3.302$ |
| % Atypical Kv1.2 domains | <u>Interaction:</u><br>$F_{1,12} = 14.47$, $p = 0.0025$<br><u>Injury:</u><br>$F_{1,12} = 293.7$, $p < 0.0001$<br><u>Drug:</u><br>$F_{1,12} = 7.281$, $p = 0.0194$ | <u>sham veh vs TBI veh:</u><br>$p < 0.0001$<br><u>sham 4-AP vs TBI 4-AP:</u><br>$p < 0.0001$<br><u>TBI veh vs TBI 4-AP:</u><br>$p = 0.0012$ | <u>sham veh vs TBI veh:</u><br>$d = 11.136$<br><u>sham 4-AP vs TBI 4-AP:</u><br>$d = 6.312$<br><u>TBI veh vs TBI 4-AP:</u><br>$d = 5.511$ |

Abbreviations: veh: saline vehicle control; ns: non-significant ( $p > 0.05$ ). Sham veh vs sham 4-AP ns for each measure ( $p > 0.4$ or higher). Veh TBI males vs veh TBI females ns for each measure ( $p > 0.7$ or higher).

**Table 2.**
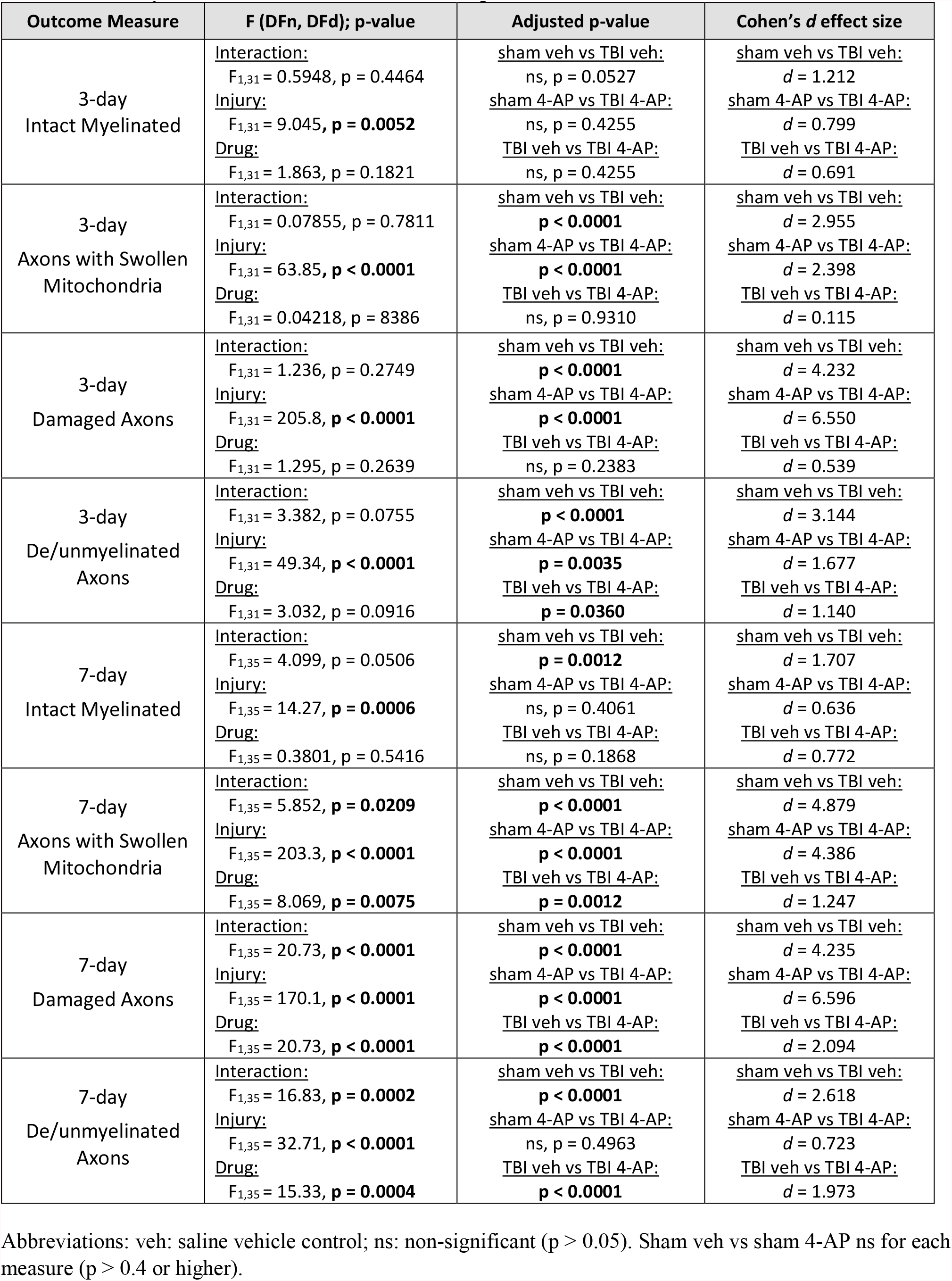
Study 2 — Electron microscopy identification of axon pathological features (Figure 4). Two -way ANOVA results with Holm-Sidak *post hoc* test and Cohen’s *d* effect size.

| Outcome Measure | F (DFn, DFd); p-value | Adjusted p-value | Cohen's *d* effect size |
| --- | --- | --- | --- |
| 3-day<br>Intact Myelinated | <u>Interaction:</u><br>$F_{1,31} = 0.5948$, $p = 0.4464$<br><u>Injury:</u><br>$F_{1,31} = 9.045$, $p = \mathbf{0.0052}$<br><u>Drug:</u><br>$F_{1,31} = 1.863$, $p = 0.1821$ | <u>sham veh vs TBI veh:</u><br>ns, $p = 0.0527$<br><u>sham 4-AP vs TBI 4-AP:</u><br>ns, $p = 0.4255$<br><u>TBI veh vs TBI 4-AP:</u><br>ns, $p = 0.4255$ | <u>sham veh vs TBI veh:</u><br>$d = 1.212$<br><u>sham 4-AP vs TBI 4-AP:</u><br>$d = 0.799$<br><u>TBI veh vs TBI 4-AP:</u><br>$d = 0.691$ |
| 3-day<br>Axons with Swollen<br>Mitochondria | <u>Interaction:</u><br>$F_{1,31} = 0.07855$, $p = 0.7811$<br><u>Injury:</u><br>$F_{1,31} = 63.85$, $p < \mathbf{0.0001}$<br><u>Drug:</u><br>$F_{1,31} = 0.04218$, $p = 0.8386$ | <u>sham veh vs TBI veh:</u><br>$p < \mathbf{0.0001}$<br><u>sham 4-AP vs TBI 4-AP:</u><br>$p < \mathbf{0.0001}$<br><u>TBI veh vs TBI 4-AP:</u><br>ns, $p = 0.9310$ | <u>sham veh vs TBI veh:</u><br>$d = 2.955$<br><u>sham 4-AP vs TBI 4-AP:</u><br>$d = 2.398$<br><u>TBI veh vs TBI 4-AP:</u><br>$d = 0.115$ |
| 3-day<br>Damaged Axons | <u>Interaction:</u><br>$F_{1,31} = 1.236$, $p = 0.2749$<br><u>Injury:</u><br>$F_{1,31} = 205.8$, $p < \mathbf{0.0001}$<br><u>Drug:</u><br>$F_{1,31} = 1.295$, $p = 0.2639$ | <u>sham veh vs TBI veh:</u><br>$p < \mathbf{0.0001}$<br><u>sham 4-AP vs TBI 4-AP:</u><br>$p < \mathbf{0.0001}$<br><u>TBI veh vs TBI 4-AP:</u><br>ns, $p = 0.2383$ | <u>sham veh vs TBI veh:</u><br>$d = 4.232$<br><u>sham 4-AP vs TBI 4-AP:</u><br>$d = 6.550$<br><u>TBI veh vs TBI 4-AP:</u><br>$d = 0.539$ |
| 3-day<br>De/unmyelinated<br>Axons | <u>Interaction:</u><br>$F_{1,31} = 3.382$, $p = 0.0755$<br><u>Injury:</u><br>$F_{1,31} = 49.34$, $p < \mathbf{0.0001}$<br><u>Drug:</u><br>$F_{1,31} = 3.032$, $p = 0.0916$ | <u>sham veh vs TBI veh:</u><br>$p < \mathbf{0.0001}$<br><u>sham 4-AP vs TBI 4-AP:</u><br>$p = \mathbf{0.0035}$<br><u>TBI veh vs TBI 4-AP:</u><br>$p = \mathbf{0.0360}$ | <u>sham veh vs TBI veh:</u><br>$d = 3.144$<br><u>sham 4-AP vs TBI 4-AP:</u><br>$d = 1.677$<br><u>TBI veh vs TBI 4-AP:</u><br>$d = 1.140$ |
| 7-day<br>Intact Myelinated | <u>Interaction:</u><br>$F_{1,35} = 4.099$, $p = 0.0506$<br><u>Injury:</u><br>$F_{1,35} = 14.27$, $p = \mathbf{0.0006}$<br><u>Drug:</u><br>$F_{1,35} = 0.3801$, $p = 0.5416$ | <u>sham veh vs TBI veh:</u><br>$p = \mathbf{0.0012}$<br><u>sham 4-AP vs TBI 4-AP:</u><br>ns, $p = 0.4061$<br><u>TBI veh vs TBI 4-AP:</u><br>ns, $p = 0.1868$ | <u>sham veh vs TBI veh:</u><br>$d = 1.707$<br><u>sham 4-AP vs TBI 4-AP:</u><br>$d = 0.636$<br><u>TBI veh vs TBI 4-AP:</u><br>$d = 0.772$ |
| 7-day<br>Axons with Swollen<br>Mitochondria | <u>Interaction:</u><br>$F_{1,35} = 5.852$, $p = \mathbf{0.0209}$<br><u>Injury:</u><br>$F_{1,35} = 203.3$, $p < \mathbf{0.0001}$<br><u>Drug:</u><br>$F_{1,35} = 8.069$, $p = \mathbf{0.0075}$ | <u>sham veh vs TBI veh:</u><br>$p < \mathbf{0.0001}$<br><u>sham 4-AP vs TBI 4-AP:</u><br>$p < \mathbf{0.0001}$<br><u>TBI veh vs TBI 4-AP:</u><br>$p = \mathbf{0.0012}$ | <u>sham veh vs TBI veh:</u><br>$d = 4.879$<br><u>sham 4-AP vs TBI 4-AP:</u><br>$d = 4.386$<br><u>TBI veh vs TBI 4-AP:</u><br>$d = 1.247$ |
| 7-day<br>Damaged Axons | <u>Interaction:</u><br>$F_{1,35} = 20.73$, $p < \mathbf{0.0001}$<br><u>Injury:</u><br>$F_{1,35} = 170.1$, $p < \mathbf{0.0001}$<br><u>Drug:</u><br>$F_{1,35} = 20.73$, $p < \mathbf{0.0001}$ | <u>sham veh vs TBI veh:</u><br>$p < \mathbf{0.0001}$<br><u>sham 4-AP vs TBI 4-AP:</u><br>$p < \mathbf{0.0001}$<br><u>TBI veh vs TBI 4-AP:</u><br>$p < \mathbf{0.0001}$ | <u>sham veh vs TBI veh:</u><br>$d = 4.235$<br><u>sham 4-AP vs TBI 4-AP:</u><br>$d = 6.596$<br><u>TBI veh vs TBI 4-AP:</u><br>$d = 2.094$ |
| 7-day<br>De/unmyelinated<br>Axons | <u>Interaction:</u><br>$F_{1,35} = 16.83$, $p = \mathbf{0.0002}$<br><u>Injury:</u><br>$F_{1,35} = 32.71$, $p < \mathbf{0.0001}$<br><u>Drug:</u><br>$F_{1,35} = 15.33$, $p = \mathbf{0.0004}$ | <u>sham veh vs TBI veh:</u><br>$p < \mathbf{0.0001}$<br><u>sham 4-AP vs TBI 4-AP:</u><br>ns, $p = 0.4963$<br><u>TBI veh vs TBI 4-AP:</u><br>$p < \mathbf{0.0001}$ | <u>sham veh vs TBI veh:</u><br>$d = 2.618$<br><u>sham 4-AP vs TBI 4-AP:</u><br>$d = 0.723$<br><u>TBI veh vs TBI 4-AP:</u><br>$d = 1.973$ |

Abbreviations: veh: saline vehicle control; ns: non-significant ( $p > 0.05$ ). Sham veh vs sham 4-AP ns for each measure ( $p > 0.4$ or higher).

**Table 3.** Study 2 — Electron microscopy analysis of fiber morphometry (Figure 5) Two -way ANOVA results with Holm-Sidak *post hoc* test and Cohen’s *d* effect size.

| Outcome Measure | F (DFn, DFd); p-value | Adjusted p-value | Cohen's d effect size |
| --- | --- | --- | --- |
| Axon Diameter | <u>Interaction:</u><br>$F_{1,28} = 0.1513, p = 0.7002$ | <u>sham veh vs TBI veh:</u><br>ns, $p = 0.1343$ | <u>sham veh vs TBI veh:</u><br>$d = 0.959$ |
| | <u>Injury:</u><br>$F_{1,28} = 11.08, p = 0.0025$ | <u>sham 4-AP vs TBI 4-AP:</u><br>ns, $p = 0.0798$ | <u>sham 4-AP vs TBI 4-AP:</u><br>$d = 1.380$ |
| | <u>Drug:</u><br>$F_{1,28} = 0.01632, p = 0.8993$ | <u>TBI veh vs TBI 4-AP:</u><br>ns, $p = 0.9287$ | <u>TBI veh vs TBI 4-AP:</u><br>$d = 0.115$ |
| Myelin Thickness | <u>Interaction:</u><br>$F_{1,28} = 1.159, p = 0.2908$ | <u>sham veh vs TBI veh:</u><br>**$p < 0.0001$** | <u>sham veh vs TBI veh:</u><br>$d = 3.081$ |
| | <u>Injury:</u><br>$F_{1,28} = 106.7, p < 0.0001$ | <u>sham 4-AP vs TBI 4-AP:</u><br>**$p < 0.0001$** | <u>sham 4-AP vs TBI 4-AP:</u><br>$d = 4.219$ |
| | <u>Drug:</u><br>$F_{1,28} = 0.09095, p = 0.7652$ | <u>TBI veh vs TBI 4-AP:</u><br>ns, $p = 0.5189$ | <u>TBI veh vs TBI 4-AP:</u><br>$d = 0.563$ |

Abbreviations: veh: saline vehicle control; ns: non-significant ( $p > 0.05$ ). Sham veh vs sham 4-AP ns for each measure ( $p > 0.6$ or higher).

**Table 4.** Study 3 — Electrophysiology of compound action potential (CAP) waveforms (Figure 6) Two-way ANOVA results with Holm-Sidak *post hoc* test and Cohen’s *d* effect size.

| Outcome Measure | F (DFn, DFd); p-value | Adjusted p-value | Cohen's *d* effect size |
| --- | --- | --- | --- |
| N1 Conduction Velocity | <u>Interaction:</u><br>$F_{1,43} = 0.05412$, $p = 0.8171$<br><u>Injury:</u><br>$F_{1,43} = 26.20$, **$p < 0.0001$**<br><u>Drug:</u><br>$F_{1,43} = 1.056$, $p = 0.3098$ | <u>sham veh vs TBI veh:</u><br>**$p = 0.0067$**<br><u>sham 4-AP vs TBI 4-AP:</u><br>**$p = 0.0016$**<br><u>TBI veh vs TBI 4-AP:</u><br>$ns$, $p = 0.6182$ | <u>sham veh vs TBI veh:</u><br>$d = 1.646$<br><u>sham 4-AP vs TBI 4-AP:</u><br>$d = 1.405$<br><u>TBI veh vs TBI 4-AP:</u><br>$d = 0.286$ |
| N2 Conduction Velocity | <u>Interaction:</u><br>$F_{1,43} = 0.2463$, $p = 0.6222$<br><u>Injury:</u><br>$F_{1,43} = 6.691$, **$p = 0.0132$**<br><u>Drug:</u><br>$F_{1,43} = 0.9846$, $p = 0.3266$ | <u>sham veh vs TBI veh:</u><br>$ns$, $p = 0.1860$<br><u>sham 4-AP vs TBI 4-AP:</u><br>$ns$, $p = 0.4377$<br><u>TBI veh vs TBI 4-AP:</u><br>$ns$, $p = 0.5970$ | <u>sham veh vs TBI veh:</u><br>$d = 0.945$<br><u>sham 4-AP vs TBI 4-AP:</u><br>$d = 0.582$<br><u>TBI veh vs TBI 4-AP:</u><br>$d = 0.577$ |
| N2 Width at Half-Maximum | <u>Interaction:</u><br>$F_{1,43} = 0.1562$, $p = 0.6947$<br><u>Injury:</u><br>$F_{1,43} = 7.827$, **$p = 0.0077$**<br><u>Drug:</u><br>$F_{1,43} = 0.1402$, $p = 0.7099$ | <u>sham veh vs TBI veh:</u><br>$ns$, $p = 0.1601$<br><u>sham 4-AP vs TBI 4-AP:</u><br>$ns$, $p = 0.3032$<br><u>TBI veh vs TBI 4-AP:</u><br>$ns$, $p = 0.9883$ | <u>sham veh vs TBI veh:</u><br>$d = 1.003$<br><u>sham 4-AP vs TBI 4-AP:</u><br>$d = 0.675$<br><u>TBI veh vs TBI 4-AP:</u><br>$d = 0.008$ |
| N2 Peak to Recovery | <u>Interaction:</u><br>$F_{1,43} = 2.797$, $p = 0.1017$<br><u>Injury:</u><br>$F_{1,43} = 5.211$, **$p = 0.0275$**<br><u>Drug:</u><br>$F_{1,43} = 0.08605$, $p = 0.7707$ | <u>sham veh vs TBI veh:</u><br>$ns$, $p = 0.0560$<br><u>sham 4-AP vs TBI 4-AP:</u><br>$ns$, $p = 0.6579$<br><u>TBI veh vs TBI 4-AP:</u><br>$ns$, $p = 0.5517$ | <u>sham veh vs TBI veh:</u><br>$d = 1.240$<br><u>sham 4-AP vs TBI 4-AP:</u><br>$d = 0.171$<br><u>TBI veh vs TBI 4-AP:</u><br>$d = 0.365$ |

Abbreviations: veh: saline vehicle control; ns: non-significant ( $p > 0.05$ ). Sham veh vs sham 4-AP ns for each measure ( $p > 0.5$ or higher).

### Study Exclusion Criteria

Predetermined exclusion criteria included: (a) 10% body weight loss at any point during experimentation (n= 0); (b) mice exhibiting an incongruous righting reflex (< 2 min for TBI animals; > 2 min for sham), a fractured, depressed skull or bleeding after impact, and/or signs of pain or distress at any point during the experiment (n= 2, fractured skull, Study 2 – Figure 1A); (c) failure to meet technical quality standards for brain slices (n= 4, anatomical region not intact after cutting). In addition, one mouse from Study 3 – Electrophysiology had to be excluded due to building site closure in response to the COVID-19 pandemic.

## RESULTS

### Characterization of study group variables and 4-AP serum levels

The effect of 4-AP on white matter injury in the acute phase of TBI was examined in the CC using a well characterized closed head model (Mierzwa et al., 2015; Marion et al., 2018; Bradshaw et al., 2021b). A pre-determined randomized, controlled study design, with blinding, was used to allocate mice to groups in each of three separate studies (Figure 1A). To evaluate comparability among the allocated groups, potential confounding variables were analyzed across the experiments. TBI mice exhibited a significantly longer righting time compared to sham controls (Figure 1B), which confirms the injury effect on this surrogate measure of altered consciousness. The similar delay in the righting reflex of TBI mice allocated to vehicle and 4-AP groups indicates comparable response to injury before the start of treatment (Figure 1B). The mean body weight among all groups of mice remained stable relative to pre-surgical measurements (Figure 1C), which shows the TBI and/or 4-AP treatment did not have detrimental effects on overall health. Furthermore, mice did not exhibit overt signs of adverse side effects such as altered grooming and posture, hyperactivity, tremors, convulsions, or cardiac arrest. Based on these observations, the low clinically relevant 4-AP dose of 0.5 mg/kg used in this study did not produce poor health or behavioral evidence of convulsant effects. 4-AP acts as a convulsant in adult mice at much higher bolus doses of over 5 mg/kg (ED_50_ of 10.9 mg/kg; (Yamaguchi and Rogawski, 1992).

Treatment with 4-AP began 24 hrs after the sham or TBI surgeries in order to provide a clinically reasonable time to initiation of treatment, as in previous studies of 4-AP treatment on acute peripheral nerve injury (Tseng et al., 2016). To determine the 4-AP circulating levels following the 0.5 mg/kg bolus injections, 4-AP was measured in cardiac serum samples drawn prior to perfusion at approximately 30-60 minutes after the last 4-AP or vehicle injection (Figure 1D). Separate experiments in a cohort of 12 mice (n = 3/group) had delayed blood sample collection (2.75 to 7.25 hours) after the last injection and found that the 4-AP levels in serum dropped to below detectable levels (data not shown), in agreement with a short (∼2 hrs) half-life in rodents (Capacio et al., 1996; Sindhurakar et al., 2017). This data demonstrates that the low 4-AP dosage used in this investigation was near or below the clinically acceptable serum dose range of 20-100 ng/ml and overt signs of unacceptable adverse side effects were not observed.

### Acute 4-AP treatment reduces axon damage in the CC after TBI

To determine the effect of 4-AP administration on axon damage after TBI, we developed a semi-automated screening assay using Thy1-YFP-16 reporter mice (Study 1, Figure 1A). In these mice, yellow fluorescence protein (YFP) is expressed in a broad spectrum of projection neurons crossing through the CC (Feng et al., 2000). In healthy adult mice, fluorescence is evenly distributed throughout the cytoplasm of axons expressing the YFP transgene. TBI produced dispersed but widespread axonal injury that was visible as robust YFP accumulations in axonal swellings throughout the CC of TBI mice in the vehicle group (Figure 2A). Only nominal levels of small YFP+ axonal accumulations were detected in sham mice (data not shown, and as published in (Marion et al., 2018). Acute 4-AP treatment on days 1-7 post-TBI dramatically reduced axonal YFP+ swellings and accumulations in TBI mice (Figure 2B).

**Figure 2.**
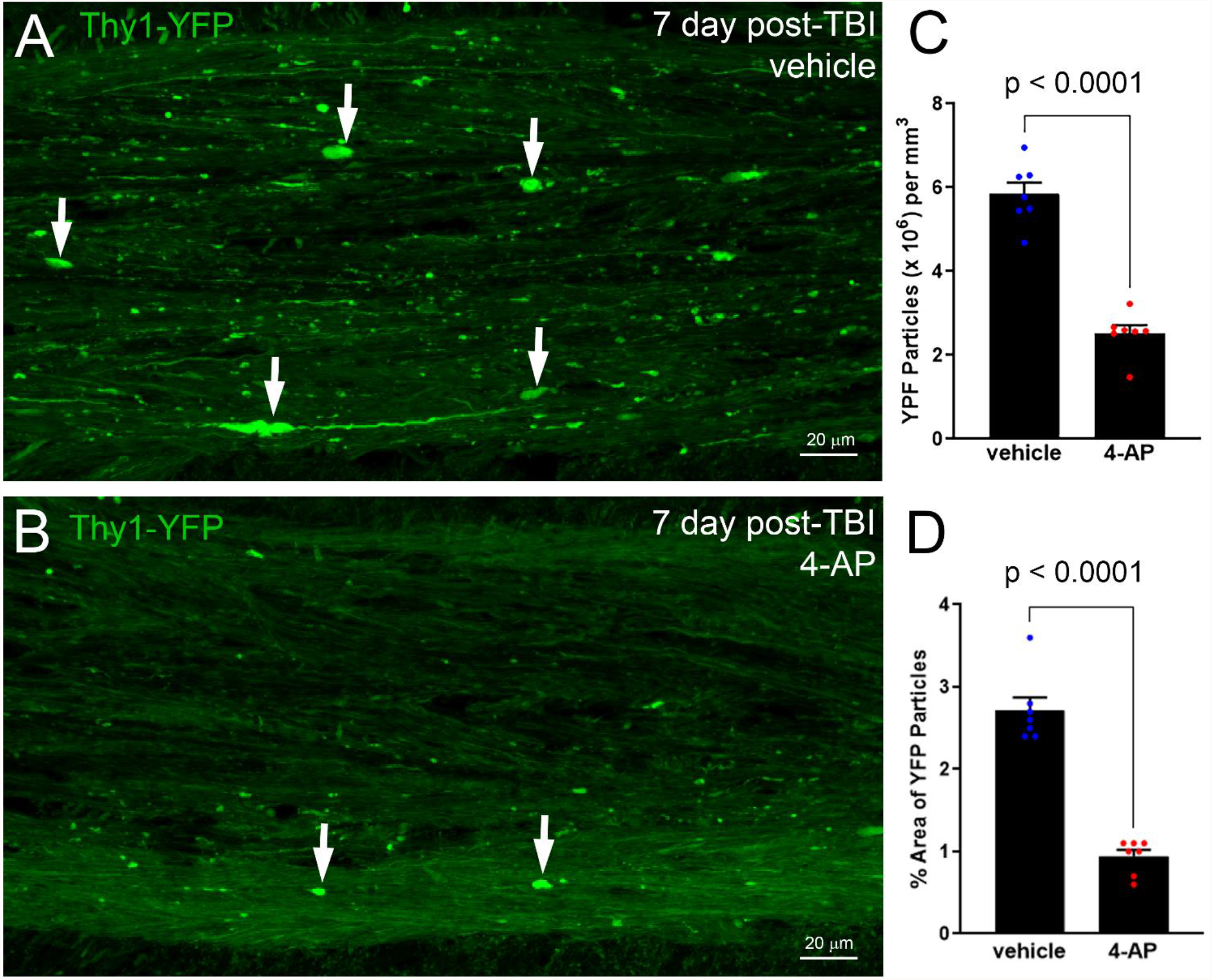
4-AP treatment shows potent axon protection using YFP-filled swellings to screen for damaged axons. *A-B:* Thy1-YFP-16 mice were given vehicle (A) or 4-AP (B) treatment on days 1-7 after TBI. Confocal images of axons in the corpus callosum illustrate YFP (green) fluorescence accumulation in axonal swellings (white arrows), which is a sensitive indicator of axon damage. Adjacent YFP-positive axons have YFP diffusely distributed along normal appearing axons. *C-D*: ImageJ particle analysis of YFP accumulation in axonal swellings. 4-AP treatment reduced the density (C) and the percent area occupied by axon swellings (D). Bars represent mean ± SEM with an individual data point shown for each mouse. Unpaired Student’s *t*-tests. See Table 1 for statistical details.

Matched regions of the CC were quantified at 7 days after TBI in mice treated with either 4-AP or vehicle, and indicated that 4-AP treatment mitigated axonal damage as revealed by YFP detection. This semi-automated approach enabled sampling of a relatively large region of the CC and takes advantage of YFP detection of subtle damage in swellings through to formation of terminal end bulbs at points of axon disconnection (Gu et al., 2017; Weber et al., 2019; Bradshaw et al., 2021a). 4-AP treatment significantly decreased the number of YFP-filled swellings (Figure 2C). The overall proportion of the CC area occupied by YFP+ axonal swellings was also significantly less in 4-AP treated mice (Figure 2D). A statistically significant difference was not found between male and female mice of either treatment group (Table 1). Therefore, subsequent studies used only male mice. Taken together, these results yielded large effect sizes for axon damage reduction following 4-AP treatment with YFP accumulations quantified based on the number (*d* = 5.231) or percent area occupied (*d* = 5.370). This screening experiment provided compelling support for more extensive analyses with continuing to initiate treatment one day after TBI and delivering 4-AP as twice daily (0.5 mg/kg) injections.

### Acute 4-AP treatment reduces disorganization of nodal regions after TBI

Axon domains associated with the node of Ranvier that are essential for rapid conduction of nerve impulses were investigated in Thy1-YFP-16 mice to explore potential 4-AP effects related to this aspect of axonal damage (Figure 3). We have previously shown TBI disrupts axonal paranodes that flank nodes of Ranvier (Marion et al., 2018). Coronal brain sections through the CC were immunolabeled for contactin-associated protein (Caspr) and the 1.6 isoform of voltage-gated sodium channels (Nav1.6). In myelinated axons, Caspr proteins are localized in axonal paranodes, which are sites of myelin attachment that flank nodes of Ranvier containing Nav1.6 channels (Figure 3A). TBI disrupted axon-myelin interactions at the paranodes based on the altered distribution of Caspr immunoreactivity at paranodes (Figure 3A). While most paired Caspr domains appear symmetrical, i.e. similar in length on both sides of a given node, others were asymmetrical (Figure 3B). Heminodes were also present as single (unpaired) Caspr immunoreactive domains adjacent to Nav1.6 labeled nodes along YFP-expressing axons (Figure 3B).

**Figure 3.**
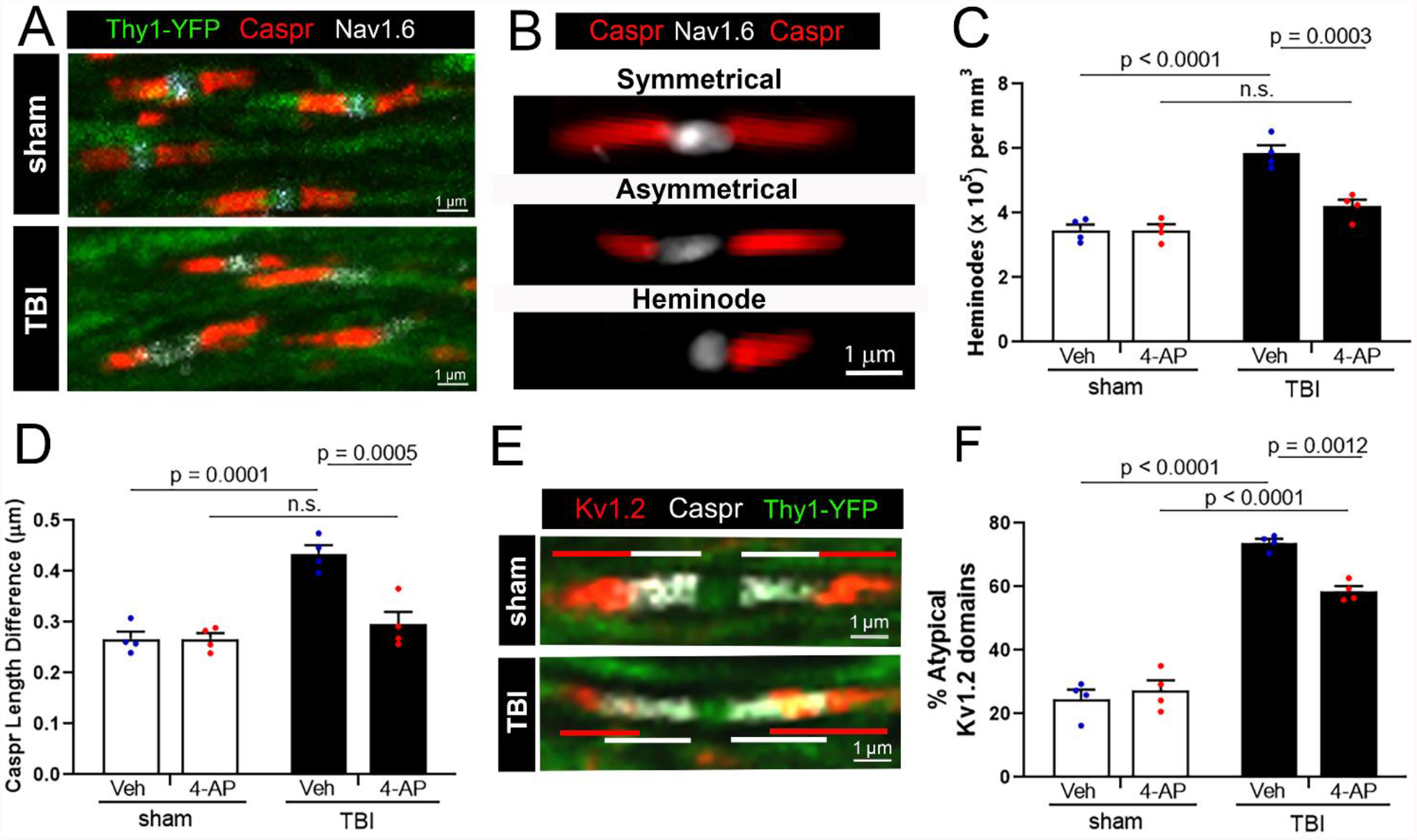
4-AP treatment improves the molecular organization of excitable axonal domains that are disrupted by TBI. Confocal imaging analysis of node of Ranvier complexes in individual corpus callosum axons of Thy1-YFP-16 mice at 7 days post-TBI or sham procedures. (A) Immunostaining along YFP-labeled axons (green) detected clustering of voltage-gated sodium channels (Nav1.6, white) at the node and Caspr (red) in the paranode region, where myelin attaches to the axon. *B-D*: TBI disrupted paranode domain organization, which was normalized by 4-AP treatment. (B) “Symmetrical” Caspr paranodes exhibit paired Caspr bands of approximately symmetrical length, while “Asymmetrical” paranodes exhibit uneven Caspr domain lengths. Single Caspr paranodes with a missing domain counterpart on the opposite side of a node were classified as “Heminodes”. (C) TBI increased the number of Caspr heminodes while acute 4-AP treatment post-TBI resulted in heminode numbers similar to sham levels. (D) TBI increased the asymmetry of Caspr domains in TBI mice compared to sham. 4-AP treatment significantly improved paranode organization following TBI. *E-F:* Kv1.2 channel (red) and Caspr (white) immunostaining along individual YFP (green) axons in single confocal optical slices. (E) In injured mice, Kv1.2 channels mislocalize from the juxtaparanode domain into Caspr-labeled paranode domain. (F) Quantification of atypical Kv1.2 domains that overlapped with Caspr domains and/or were asymmetrical in length. Acute 4-AP treatment reduced the percentage of atypical Kv1.2 domains after TBI, but the distribution patterns of Kv1.2 channels was not fully normalized to sham levels. *C, D, F*: Bars represent mean ± SEM with an individual data point shown for each mouse. Two-way ANOVA with Holm-Sidak’s multiple comparisons test. See Table 1 for statistical details.

We found that 4-AP treatment protected against TBI-induced changes in Caspr paranode domains. In the vehicle conditions, TBI increased Caspr heminodes and asymmetry in the paranodes as compared to sham mice (Figure 3C, D). 4-AP treatment led to a substantial decrease in post-TBI heminodes and asymmetrical paranodes, with levels comparable to sham mice at the 7-day study endpoint (Figure 3C, D). The nodal gap between Caspr pairs was not significantly different between groups (data not shown).

Immunolabeling Thy1-YFP-16 sections for Kv1.2 channel to examine channel distribution revealed that 4-AP treatment significantly improved Kv1.2 localization. Axonal Kv1.2 voltage-gated potassium channels, which are a potential therapeutic target of 4-AP, are localized in juxtaparanode regions under the myelin sheath in healthy myelinated axons. Caspr containing paranodes segregate the ionic currents of potassium channels in juxtaparanodes from the sodium channels in the node of Ranvier, which is critical for action potential propagation. In healthy axons, Caspr paranode domains and adjacent Kv1.2 juxtaparanode domains are clearly separated with minimal overlap (see sham example in Figure 3E). TBI caused a loss of domain boundaries, with Caspr and Kv1.2 domains markedly overlapping with each other (see TBI example in Figure 3E). In vehicle controls, TBI caused disordered atypical Kv1.2 immunolabeling patterns (Figure 3F). Specifically, TBI increased the proportion of axons with Kv1.2 overlapping into paranodal (Caspr+) regions, asymmetry between paired Kv1.2 domains, and/or Kv1.2 dispersed away from the nodal region. After TBI, 4-AP treatment significantly improved Kv1.2 localization as compared to the vehicle condition (Figure 3F). However, atypical Kv1.2 domains in TBI mice remained significantly above sham levels. This analysis shows significant improvement, but not full restoration, of the complex molecular domains of nodal regions during the first week post-TBI with 4-AP treatment.

### Acute 4-AP treatment reduces axon damage and myelin loss during the first week post-TBI

While YFP fluorescence sensitively detects axon swellings as an indicator of damage, electron microscopy is the gold standard approach to examine distinct features of pathology in axons and their ensheathing myelin. Electron microscopy was used to evaluate 4-AP effects on axon damage and demyelination in the CC after TBI (Study 2, Figure 1A). Studies were conducted with 4-AP (0.5 mg/kg; i.p.) or saline vehicle injections initiated at 24 hours after TBI or sham procedures and continued twice daily until perfusion on day 3 or 7. The 3-day time point was chosen to allow comparison with our previous ultrastructural characterization of axon and myelin pathology in this concussive TBI model (Mierzwa et al., 2015; Marion et al., 2019). The 7-day time point aligns with the analysis in Thy1-YFP-16 mice for in-depth interpretation of the axon damage and nodal region pathology (Figures 2, 3).

Quantitative analysis showed that TBI reduced the healthy-appearing, i.e. intact, myelinated axons (Figure 4A). In agreement with this finding, in the vehicle condition, TBI resulted in significant pathology in myelinated axons that was identified based on specific ultrastructural features. Healthy-appearing axons exhibiting only swollen mitochondrial profiles were increased after TBI (Figure 4B) and may represent an early, and possibly reversible, state of axonal injury (Balan et al., 2013). TBI resulted in damaged axons that exhibited condensed cytoplasm, with compacted cytoskeletal structure, and vesicle or organelle accumulations (Figure 4C). Axons with diameters greater than 0.3 µm, which are typically large enough to be myelinated in adult mouse CC, that lacked myelin were classified as de/unmyelinated axons. TBI-induced demyelination was indicated by de/unmyelinated values above sham levels (Figure 4D).

**Figure 4.**
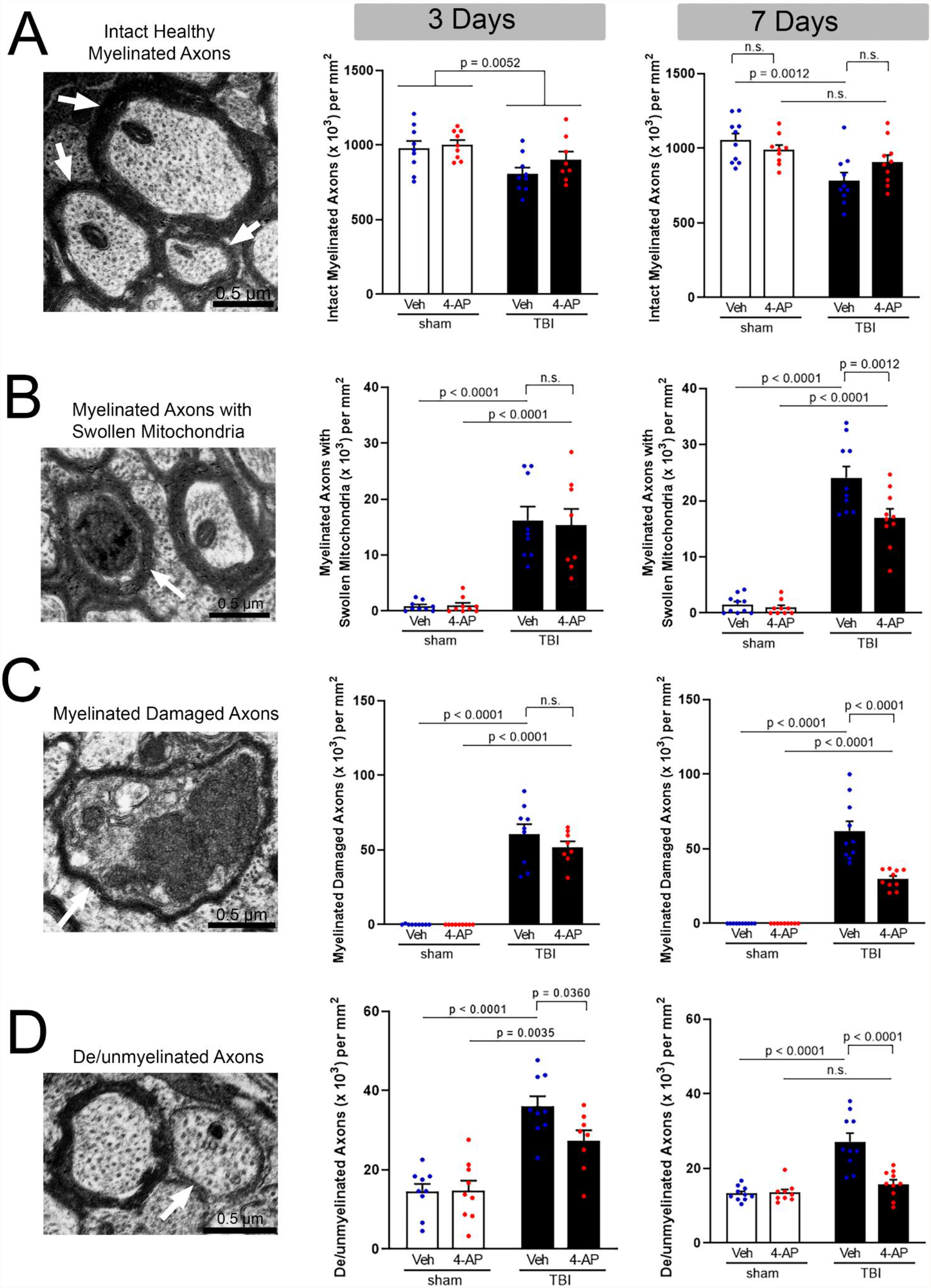
4-AP benefit requires continued treatment during the first week post-TBI to protect against multiple features of axon damage. Electron microscopy imaging of axon and myelin ultrastructural features (left) with classification of axon pathology at 3 days (middle) or 7 days (right) after TBI or sham procedure. *Left panels:* Examples of axon and myelin ultrastructural features. (A) Intact healthy myelinated axons (arrows). (B) Swollen mitochondrion (i.e., occupying over 50% of the axon cross section; arrow) in contrast to the typical mitochondrion in the adjacent axon. (C) Damaged myelinated axon (arrow) exhibiting condensed cytoplasm and vesicle/organelle accumulation. (D) Demyelinated axon (arrow) lacking a myelin sheath but structurally intact with diameter > 0.3 µm, which is typically myelinated. *Middle Panels:* At the 3-day time point, TBI significantly reduced the number of intact myelinated axons (A), and increased myelinated axons with swollen mitochondria (B) or axon damage (C), and induced demyelination of additional axons (D). 4-AP treatment on days 1-3 did not have a significant benefit as compared to vehicle. (B) or the number of damaged (C) and demyelinated (D) axons post-TBI. *Right Panels:* With 4-AP treatment extended to 7 days, the number of intact myelinated axons post-TBI was statistically similar to sham levels (A). More specifically, 4-AP treatment significantly reduced axons exhibiting mitochondrial swelling (B), axon damage (C), or TBI-induced demyelination (D). *A-D*: Bars represent mean ± SEM with an individual data point shown for each mouse. Two-way ANOVA with Holm-Sidak’s multiple comparisons test. See Table 2 for statistical details.

4-AP treatment through 7 days showed significant benefits on axon and myelin pathology, which were not evident at 3 days post-TBI. At the 7 day time point, as compared to vehicle, 4-AP significantly reduced TBI pathology for axons with swollen mitochondria by 29.5% (Figure 4B), damaged axons by 51.7% (Figure 4C), and de/unmyelinated axons by 41.7% (Figure 4D). In the case of axons with swollen mitochondria, 4-AP treatment appeared to attenuate further deterioration in the axons. In the case of damaged and de/unmyelinated axons, in contrast, 4-AP treatment appeared to enhance recovery. This reduced pathology was still significantly above sham levels for mitochondrial swelling and axon damage while de/unmyelinated axon values recovered to sham levels. These results demonstrate that 4-AP reduces axon damage and demyelination during a therapeutic window beginning one day after the mechanical injury and continuing through the first week post-TBI.

The EM images of CC axons were further examined to quantify axon diameter and myelin thickness, which can both influence axon conduction properties (Figure 5). The mean diameter of myelinated axons decreased significantly in TBI conditions compared to sham mice (Figure 5A). TBI also significantly reduced the myelin sheath thickness in both vehicle and 4-AP treated mice compared to sham controls (Figure 5B). A specific effect of 4-AP treatment was not found for either axon diameter or myelin thickness. In adult white matter, myelin thickness varies with axon diameter. This relationship was examined using the g-ratio, which is the ratio of the axon diameter to the diameter of the myelinated axon fiber. Comparison of the g-ratio relative to axon diameter showed that myelin was thinner due to TBI as compared to sham, which was found in both 4-AP and vehicle groups (Figure 5C, D). The g-ratio was not changed by 4-AP treatment in sham or TBI groups (Figure 5E, F).

**Figure 5.**
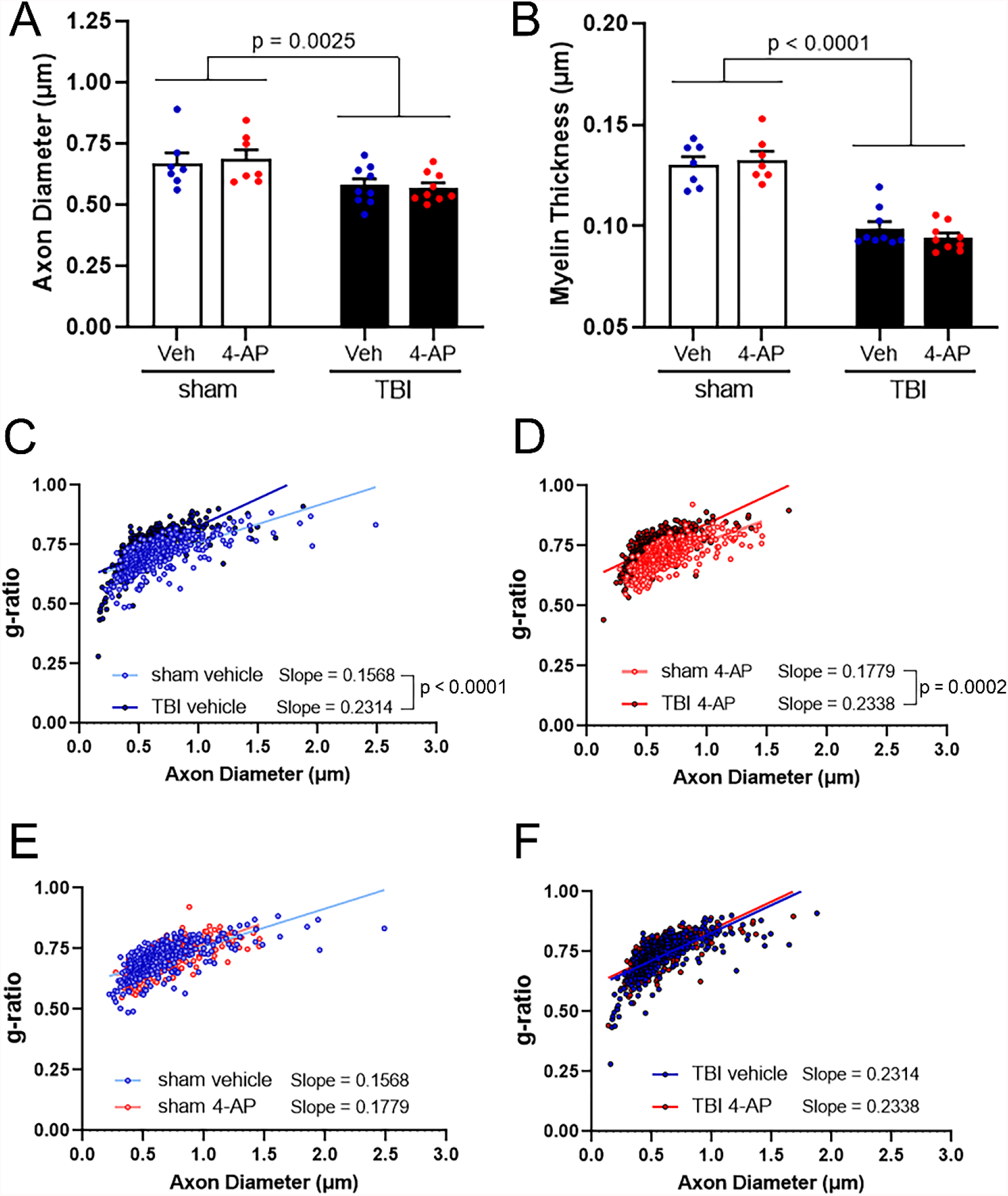
TBI reduces axon diameter and myelin thickness in 4-AP and vehicle conditions. Morphological analysis of intact myelinated axons in the corpus callosum of mice treated with 4-AP or vehicle on days 1-7 post-TBI or sham condition. (A) TBI results in atrophic axons in both vehicle and 4-AP treated mice. (B) Intact axons have thinner myelin after TBI in both vehicle and 4-AP treated mice. *C-F:* Scatter plots display axon diameter against g-ratio (inner axonal diameter divided by total outer diameter), since appropriate myelin thickness is related to the diameter of a given axon. The slope of the linear regression was steeper for TBI versus sham mice. This indicates myelin thinning in proportion to axon diameter after TBI. This relationship was not influenced by 4-AP treatment as compared to vehicle (E, sham p = 0.1362; F, TBI p = 0.8818). *A-B*: Two-way ANOVA with Holm-Sidak’s multiple comparisons test Bars represent mean ± SEM with an individual data point shown for each mouse. *C-F*: Linear regression analysis. Each circle represents a measured axon (50 axons/mouse). See Table 3 for statistical details.

### Acute 4-AP treatment does not ameliorate functional deficits of CC axon populations within the first week after TBI

To evaluate axon function relative to TBI and systemic 4-AP administration, *ex vivo* studies of brain slices were used to directly measure signal conduction properties of CC axons (Study 3, Figure 1A). Studies were conducted with 4-AP (0.5 mg/kg; i.p.) or saline vehicle injections initiated at 24 hours after TBI or sham procedures and continued twice daily through day 6. Brains were collected for electrophysiology on day 7 after an overnight period to allow for washout of circulating 4-AP. The compound action potentials (CAPs) recorded using graded current intensities reflect the summed response of a population of axons successfully conducting from the stimulus electrode across the CC midline to reach the recording electrode (Figure 6A). In the CC, CAPs form two distinct wave peaks based on the conduction velocity (Figure 6B, C). As shown in the representative traces (Figure 6C, the first wave to reach the recording electrode (N1) represents the response of fast-conducting myelinated axons, while the second wave (N2) comprises the response of slower conducting non-myelinated axons. TBI slowed conduction velocities in both the N1 and N2 axon populations (Figure 6B). Acute 4-AP treatment did not alter conduction velocity in sham or TBI mice, as compared to the vehicle condition. The amplitude of the N1 CAP field potential appeared noticeably diminished in recordings from TBI animals compared to sham controls (Figure 6C). Plotting the CAP amplitude against stimulus intensity revealed that the N1 CAP amplitude was consistently lower in mice that received a TBI as compared to the sham controls (Figure 6 D). The N2 CAP amplitude was not altered by TBI or 4-AP treatment (Figure 6D). However, TBI did affect the shape of the N2 wave resulting in a widened waveform that recovered more slowly (Figure 6E-G), which reflects a broader range of axon velocities particularly those with slower conduction velocity. The shape of the N1 wave was not altered by TBI or 4-AP (data not shown).

**Figure 6.**
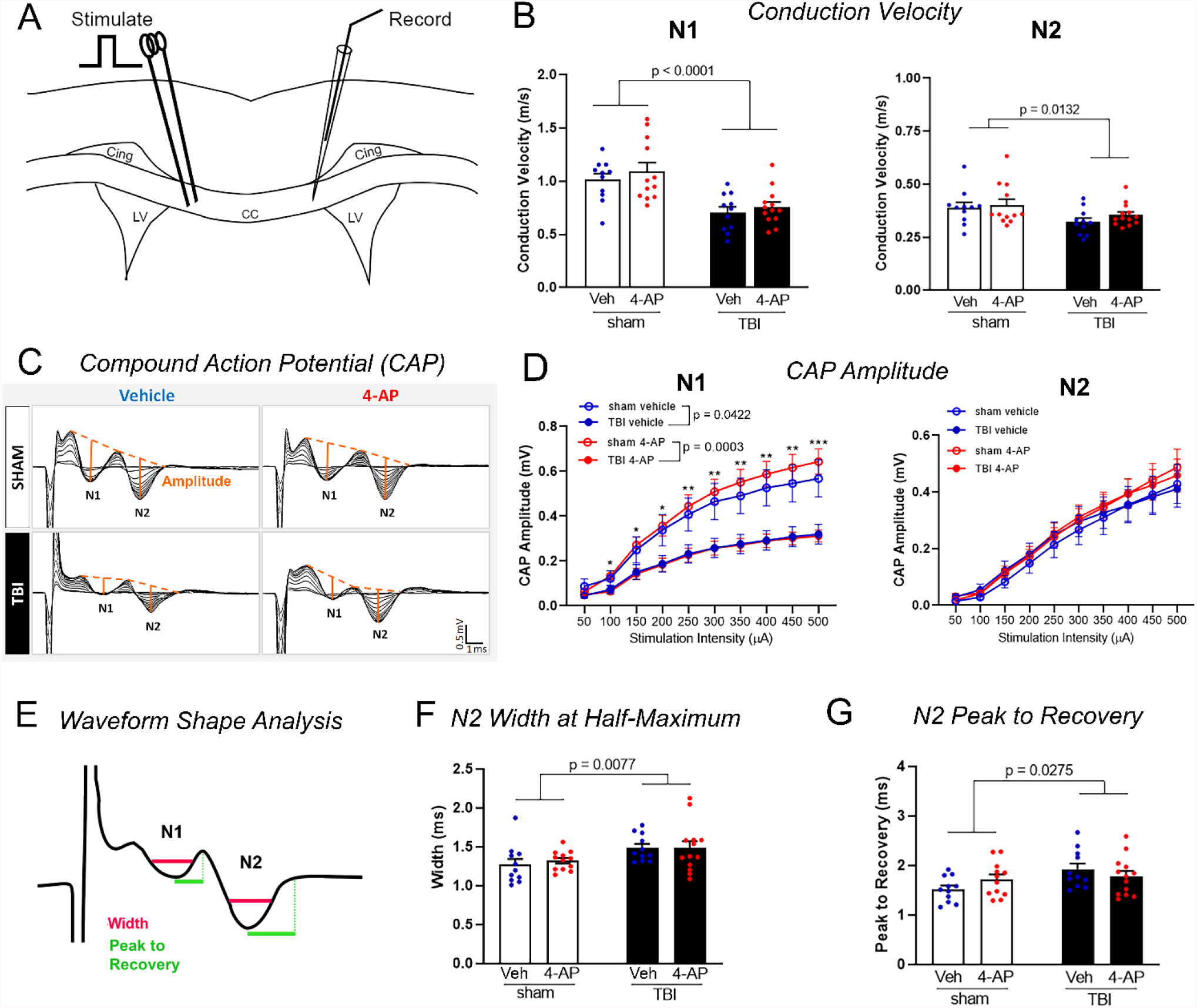
TBI slows the axon compound action potential velocity and amplitude. Electrophysiology recordings were used to directly test the function of axons in the corpus callosum. (A) Schematic of the position of the stimulating and recording electrodes for measurements of axon compound action potentials (CAP) and conduction velocity in *ex vivo* brain slices. (B) TBI reduced the speed of action potential conduction in both faster N1 wave comprised of myelinated axons and slower N2 wave comprised of unmyelinated and potentially demyelinated axons. The 7-day 4-AP treatment regimen did not restore this velocity deficit. (C) Representative input-output traces show the evoked N1 and N2 CAP waveforms with stimulus intensity pulses ranging from 50 to 500 µA (50 µA increments). Orange lines from CAP peaks to their projected bases indicate the amplitude of the response of fibers at a given stimulus intensity. (D) CAP amplitude analysis revealed a main injury effect on myelinated axons based on the reduced amplitude of only the N1 wave. N1 CAP amplitude difference was significant with post-hoc comparison for the sham versus TBI mice with 4-AP treatment. (E) Schematic of complementary spike waveform parameters. The width is dependent on the conduction velocity distribution among contributing axons. The time from peak to recovery represents the time for membrane repolarization among the slowest conducting axons comprising each waveform. The CAP width (F) and time from peak to recovery (G) indicated that TBI prolonged the recovery of the N2 waveform in both vehicle and 4-AP conditions. The N1 wave shape parameters were not altered by TBI or 4-AP treatment (data not shown). Abbreviations: Cing, cingulum; CC, corpus callosum; LV, lateral ventricles. *B, F, G*: Bars represent mean ± SEM with individual data points for each mouse (n = 11-13 animals per group). *D*: Linear effects model statistical analysis and Holm-Sidak’s multiple comparison test (*p < 0.05, **p < 0.01, *** p < 0.001). Two-way ANOVA and Holm-Sidak’s multiple comparisons test. See Table 4 for statistical details.

Further analysis tested whether TBI and/or systemic 4-AP treatment altered the intrinsic excitability or refractory period of CC axons (Figure 7). Following CAP amplitude and velocity recordings, additional recordings examined evoked CAP strength-duration properties. The threshold for activation of nerve fibers depends not only on stimulus strength, as shown with CAP amplitude analysis, but also on the electric charge from the product of the strength and duration of the stimulus. In order to probe the relationship between minimal intensity (current) of an electrical pulse and stimulus pulse duration, strength-duration testing was conducted as previously described (Reeves et al., 2005). This analysis showed that TBI significantly altered the strength-duration characteristics of the N2 CAP waveform at shorter pulse durations without affecting the excitability of the N1 wave (Figure 7A). The leftward shift in the plot of stimulus current versus stimulus duration for sham vehicle compared to TBI vehicle animals suggests that slower conducting (N2) axons are more excitable following TBI. We also found that 4-AP treatment increased the excitability of both the N1 and N2 components of the CAP in sham mice, but not in TBI animals (Figure 7A). This result suggests that *in vivo* 4-AP exposure decreases the stimulation threshold of CC axons, so less electric charge is sufficient to bring responsive fibers to threshold. Importantly, 4-AP did not exacerbate axonal excitability properties following TBI.

**Figure 7.**
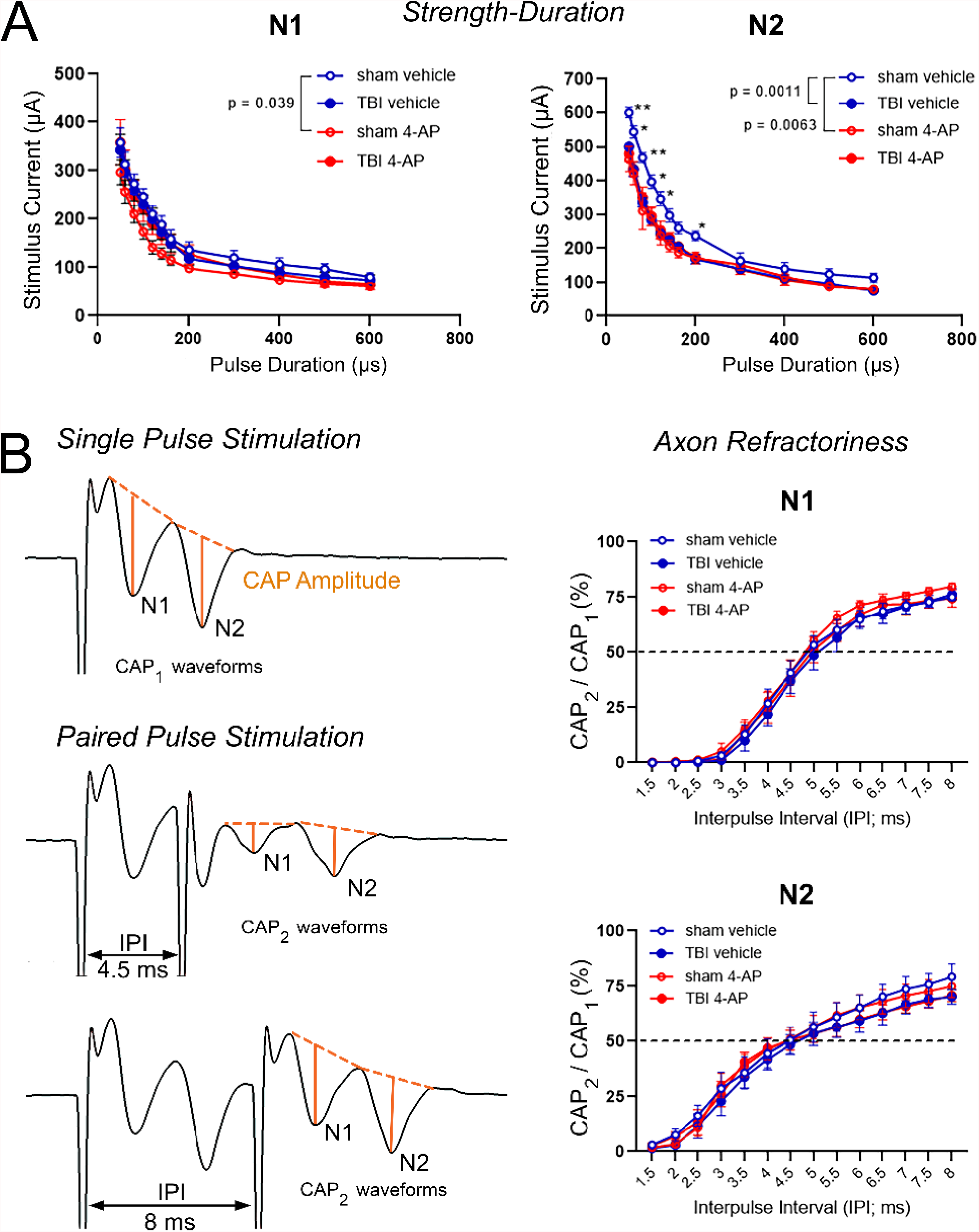
TBI alters the axon population excitability without changing the refractory period. A subset of mice had extended studies of additional conduction properties from the *ex vivo* brain slice preparations as shown in Figure 6. (A) The axon intrinsic excitability was probed by measuring the stimulus duration relative to increasing stimulus current. This strength-duration analysis revealed that TBI induced hyperexcitability of N2 axons, based on eliciting a similar response with a lower stimulus strength at a given stimulus duration. This difference in the N2 axon excitability was significant with post-hoc comparison for the sham versus TBI with vehicle. In both N1 and N2 axons, 4-AP treatment shifted the strength-duration curve toward a more excitable state in sham mice, but not in TBI mice. (B) The axon refractory period due to recovery time between action potentials was probed with a paired-pulse protocol. Representative N1 and N2 compound action potential (CAP) waveforms from a single pulse stimulation as compared to a paired-pulse protocol with varying interpulse intervals (IPI). Orange lines from CAP peaks to their projected bases indicate the amplitude of each waveform. Plots of the percent ratio of each second pulse CAP amplitude (CAP_2_), evoked in the paired-pulse stimulation, divided by the CAP amplitude of the single reference pulse stimulation (CAP_1_) showed no differences in refractoriness between sham and TBI animals or between vehicle and 4-AP treatments. *A-B*: Linear effects model statistical analysis and Holm-Sidak’s multiple comparison test (*p < 0.05, **p < 0.01). Data is expressed as mean ± SEM with n = 6-7 animals per group.

Finally, the refractory properties of CC axons were analyzed using a paired pulse paradigm in the brain slices as for strength-duration testing. After firing an action potential, axons enter a refractory period of reduced excitability. Axon refractoriness reflects the minimal interpulse interval needed to depolarize the membrane from a hyperpolarized state into a threshold capable of initiating a second action potential. Juxtaparanode Kv1 channels contribute to repolarize axons between impulses, which may be disrupted during demyelination (Rasband et al., 1998; Devaux et al., 2002; Cohen et al., 2020). To analyze axon refractoriness, the amplitudes of electrically evoked CAPs elicited by a paired pulse stimulation (CAP_2_ waveforms) were compared to the amplitude of their respective N1 and N2 CAP components from a single pulse stimulation (CAP_1_ waveforms) (Figure 7B). Neither the TBI nor 4-AP treatment affected the refractoriness of the N1 or the N2 wave components.

Taken together, our electrophysiological results indicate that the main functional deficits after TBI impact the speed and population size of myelinated axons that conduct impulses across the CC, which was not altered by acute 4-AP treatment when examined at the 7 day post-TBI.

## DISCUSSION

The current pre-clinical study is the first test of repurposing 4-AP as an acute treatment for TBI, and our results show significant benefit for treating axon damage. 4-AP treatment initiated at a clinically reasonable time of one day after a TBI was effective at a low clinically relevant dosage that produced robust beneficial effects on CC axon and myelin pathology within the first week after TBI. 4-AP significantly reduced axon damage using two distinct techniques in separate studies (Figures 1-4). Using axonal YFP accumulations as a sensitive screen of axon damage showed the most robust 4-AP effect sizes, and demonstrated similar results in both sexes (Figure 2). Significant positive 4-AP effects on axonal pathology were also identified based on mitochondrial swelling, impaired vesicle transport, and cytoskeletal compaction (Figure 4). Furthermore, 4-AP significantly reduced axonal demyelination and disruption of nodal regions after TBI (Figures 3, 4). These benefits of acute 4-AP treatment were found in a closed head TBI model that produced dispersed damaged axons distributed among intact axons within the white matter, reflecting the hallmark pathological characteristics of diffuse axonal injury in cases of human TBI (Smith et al., 2013; Smith and Stewart, 2020).

Our results also suggest that 4-AP treatment enhances repair of multiple myelination-related outcomes. At 7 days post-TBI, myelin sheaths were abnormally thin as compared to sham values in both 4-AP and vehicle cohorts (Figure 5), a feature associated with remyelination and preservation of axonal function in experimental demyelination (Duncan et al., 2017). Consistent with ongoing remyelination, demyelination decreased significantly between 3 and 7 days after TBI (p = 0.0066 vehicle; p = 0.0004 4-AP), but it was only in the 4-AP treated group that demyelination returned to sham levels (Figure 4). Furthermore, heminodes have been shown to form during developmental myelination or where remyelination is incomplete (Coman et al., 2006; Freeman et al., 2016; Brivio et al., 2017), and the increased frequency of heminodes after TBI was reduced to sham levels only in the 4-AP treated mice (Figure 3). Interpreting the 4-AP reduction of heminodes as reparative is supported by our prior study that showed increased heminode frequency occurs by at least 3 days post-TBI and does not resolve but instead further increases at 6 weeks (Marion et al., 2018). Collectively, these findings (Figures 2-5) suggest that TBI causes axon damage with myelin sheath detachment from paranodes and demyelinated axon segments, and that 4-AP treatment enhances recovery from this damage.

The outcomes of our studies provide further evidence that treatment of acute neurological injuries with 4-AP provides multiple unexpected benefits. In acute peripheral nerve crush injuries in mice, initiation of treatment with clinically relevant concentrations of 4-AP one day post-injury promoted remyelination and enhanced axonal area (Tseng et al., 2016). 4-AP treatment has also been reported to prevent onset of experimental allergic enchephalomyelitis and decrease damage from optic nerve crush in mice (Dietrich et al., 2020), but these studies initiated 4-AP one week prior to immunization or injury, with daily dosages more than 10-fold greater than those shown to provide clinically relevant serum concentrations of 4-AP in the current study (Figure 1) and in oral treatment of peripheral nerve injury (Hsu et al., 2020).

The mechanisms by which relatively short-term treatment with 4-AP, initiated 24 hours post-injury at clinically relevant dosages, enhances recovery in acute neurological injuries are unknown, and may be multi-faceted. 4-AP is thought to inhibit Kv channels but the overall result also depends on sodium and calcium channel activity, and is hard to predict at low 4-AP concentrations (Wu et al., 2009; Agren et al., 2019; Dietrich et al., 2021). Beyond potential 4-AP actions at nodal regions, low 4-AP levels may have an activity dependent effect on axon and myelin health from Kv channel inhibition that potentiates synaptic transmission to enhance axon and neural circuit activity (Smith et al., 2000; Sindhurakar et al., 2017). 4-AP has also shown symptomatic benefit to improve the function of regenerating axons (Bei et al., 2016; Liu et al., 2017). Finally, 4-AP mechanisms in TBI may involve potassium channels on multiple cell types, including neurons, glia, or immune cells, and pathological conditions change the pattern of potassium channel expression and localization (Judge and Bever, 2006; Reeves et al., 2012; Calvo et al., 2016; Jukkola et al., 2017; Boscia et al., 2021; Dietrich et al., 2021).

Despite the multiple benefits of 4-AP treatment on axon pathology in our TBI studies, the results differ from outcomes on peripheral nerve injury with respect to recovery of nerve function. However, the electrophysiological deficits of CC axons observed after TBI (Figure 6) are consistent with the structural aspects that were not altered by 4-AP treatment. The N1 axon conduction velocity is dependent on axon diameter and myelin thickness (Waxman, 1980), which were reduced after TBI (Figure 5). The lower N1 CAP amplitude indicates that fewer axons conducted signal through the CC after TBI due to loss of damaged axons or impaired conduction along viable axons (Franssen, 2008; Uncini and Santoro, 2020; Duncan et al., 2021). These electrophysiological findings are in agreement with prior studies in CC axons after TBI from our lab and others (Reeves et al., 2005; Dileonardi et al., 2012; Reeves et al., 2012; Reeves et al., 2016; Marion et al., 2018). The lack of 4-AP benefit on axon conduction properties could have been because TBI is a more complex injury than peripheral nerve crush or/and because more time is needed to observe benefits on these parameters.

The current results focus on the critical first week after TBI, when axons are highly vulnerable, and definitively show that 4-AP is acting during the acute phase to reduce axon pathology. Evaluating functional improvement with 4-AP treatment after TBI may require longer term studies and/or additional approaches. Our electrophysiological approach provided a direct measure of conduction properties among the CC axon population, but can only detect changes involving a large number of axons that achieve a capability threshold of successfully conducting action potentials across the CC. Action potential transmission may fail in surviving or recovering axons due to impedance mismatch between myelinated and incompletely remyelinated areas, or, due to current shunting at unstable axon-myelin paranode junctions (Utzschneider et al., 1994; Cohen et al., 2020), which may be factors at our 7 day time point. During remyelination, proper Kv1 localization in juxtaparanodes occurs at a late stage of node formation (Rasband et al., 1998; Zoupi et al., 2013) and Kv1.2 localization to juxtaparanodes was significantly improved with 4-AP treatment but had not recovered to sham levels (Figure 3). Interestingly, previous animal and human studies indicate functional improvement may require longer duration studies of 4-AP treatment. Low dose 4-AP treatment initiated one day after peripheral nerve crush resulted in significant gait improvement at 3 days post-injury that progressed through 8 days, although a significant increase in nerve conduction velocity was not observed until 21 days (Tseng et al., 2016). Furthermore, clinical trials of 4-AP (dalfampridine) for patients with chronic MS used a treatment duration of 4 weeks or more to evaluate improvement of function (Zhang et al., 2021).

While identifying the mechanism(s) by which 4-AP treatment decreases damage and enhances recovery after TBI or peripheral nerve injury will be challenging, it is perhaps more important that repurposing of 4-AP for examination in these traumatic injuries is more straightforward. Due to the long-standing interest in the use of 4-AP in treating multiple chronic neurological problems (including MS, nystagmus, Lambert-Easton myasthenic syndrome and spinal cord injury), extensive information exists regarding appropriate dosage regimens, side-effects and other concerns related to translation of studies from the laboratory to the clinic. Studies on diagnosis and recovery from peripheral nerve injury have already led to two clinical trials on 4-AP treatment of acute injuries (NCT03701581, NCT04026568). The need for treatments for acute TBI and the promising results of our present study suggest the importance of repurposing 4-AP for TBI as a new clinical indication.

Several limitations should be considered in interpreting these findings. Low dose 4-AP was administered twice daily, as for dalfampridine, but without the extended release achieved in the capsule formulation. This initial study focused on white matter injury, specifically CC axon and myelin pathology, whereas 4-AP effects may also involve glial cells or peripheral immune cells (Dietrich et al., 2021). Broader 4-AP effects may involve neural circuit and synaptic properties, and the fidelity of high frequency signal transmission (Yang et al., 2014; Lang-Ouellette et al., 2021; Zang and Marder, 2021), which may require more extensive electrophysiological studies.

The data from this first study of 4-AP in TBI warrant further investigation of the mechanism(s) of acute 4-AP benefit and the potential to improve longer term functional recovery. Preserving axons at the acute stage should have the greatest potential to prevent persistent symptoms. 4-AP treatment later after TBI may also attenuate neurodegeneration based on retrospective evidence in patients with MS (Dietrich et al., 2020). An effective treatment to reduce axon damage will be a tremendous benefit at any stage after TBI.

## Acknowledgements

This work was supported by the U.S. Department of Defense through the Congressionally Directed Spinal Cord Injury Research Program (SC160213) and the USU-UCSF Partnership: Brain Injury and Disease Prevention, Treatment and Research. We would like to thank Drs. Spencer Millen, Fengshan Yu, Genevieve Sullivan, and Dennis McDaniel (USUHS) for providing experimental guidance and technical assistance. We thank the USU Preclinical Behavior and Modeling Core for support of the brain injury model. We also thank Dr. Heather Natola and Ms. Kourtney Korczac (University of Rochester) for help with serum sample analyses. The opinions and assertions expressed herein are those of the author(s) and do not necessarily reflect the official policy or position of the Uniformed Services University or the Department of Defense.

